# Dual function of the O-antigen WaaL ligase of *Aggregatibacter actinomycetemcomitans*

**DOI:** 10.1101/2022.10.31.514599

**Authors:** David R. Danforth, Marcella Melloni, Richard Thorpe, Avi Cohen, Richard Voogt, Jake Tristano, Keith P. Mintz

**Affiliations:** Department of Microbiology and Molecular Genetics, University of Vermont, Burlington, VT

**Keywords:** Glycosylation, Adhesins, Autotransporter proteins, Periodontitis

## Abstract

Protein glycosylation is critical to the quaternary structure and collagen binding activity of the extracellular matrix protein adhesin A (EmaA) associated with *Aggregatibacter actinomycetemcomitans*. The glycosylation of this large, trimeric autotransporter adhesin is postulated to be mediated by WaaL, an enzyme with the canonical function to ligate the O-polysaccharide (O-PS) antigen with a terminal sugar of the lipid A-core oligosaccharide of lipopolysaccharide (LPS). In this study, we have determined that the *Escherichia coli waaL* ortholog (*rflA*) does not restore collagen binding of a *waaL* mutant strain of *A. actinomycetemcomitans* but does restore O-PS ligase activity following transformation of a plasmid expressing *waaL*. Therefore, a heterologous *E. coli* expression system was developed constituted of two independently replicating plasmids expressing either *waaL* or *emaA* of *A. actinomycetemcomitans* to directly demonstrate the necessity of ligase activity for EmaA collagen binding. Proper expression of the protein encoded by each plasmid was characterized, and the individually transformed strains did not promote collagen binding. However, co-expression of the two plasmids resulted in a strain with a significant increase in collagen binding activity and a change in the biochemical properties of the protein. These results provide additional data supporting the novel hypothesis that the WaaL ligase of *A. actinomycetemcomitans* shares a dual role as a ligase in LPS biosynthesis and is required for collagen binding activity of EmaA.

**Importance:** The human oral pathogen *A. actinomycetemcomitans* is a causative agent of periodontal and several systemic diseases. The organism expresses an adhesin, EmaA, important for the colonization of this pathobiont via collagen binding and biofilm formation. EmaA is suggested to be modified with sugars and the modification is mediated using the same enzymes involved in lipopolysaccharide (LPS) biosynthesis. In this study, evidence is presented which suggests that the WaaL ligase, the enzyme that ligates the O-polysaccharide (O-PS) antigen with a terminal sugar of the lipid A-core oligosaccharide of LPS, is required for the collagen binding activity of EmaA. This finding represents a new paradigm for the posttranslational modification of this type of autotransporter protein.

## Introduction

Gram-negative bacteria are distinguishable based on the presence of an inner and outer double membrane encircling the cytosol (1). The inner membrane is composed exclusively of phospholipids and proteins (1, 2), whereas the asymmetric outer membrane has lipopolysaccharides (LPS) replacing a majority of the phospholipids in the outer leaflet of the bilayer (2). LPS can be divided into three structural domains: Lipid A, the core oligosaccharide, and the O-polysaccharide (O-PS) (3). The O-PSs are assembled as undecaprenyl diphosphate (Und-PP)-linked assembly intermediates in the cytoplasm, transported across the inner membrane and attached to the Lipid A-core oligosaccharide by the O-antigen ligase or WaaL (4, 5). The WaaL enzyme is canonically associated with LPS biosynthesis, but studies with the oral pathogen *Aggregatibacter actinomycetemcomitans* suggest a novel dual-functionality for this enzyme (6, 7).

*A. actinomycetemcomitans* is one of a cohort of bacteria associated with periodontal disease (8, 9). Uniquely, this bacterium is associated with multiple forms of periodontal disease: localized aggressive periodontitis (LAP), an acute disease affecting children and young adults (10–14); and chronic periodontitis, a slowly developing disease affecting older individuals (10). *A. actinomycetemcomitans* also colonizes non-oral areas of the body and has been reported to be the causative agent of several diseases, including infective endocarditis, meningitis, septicemia, brain abscess and osteomyelitis (15–18). Although the oral cavity is the primary colonization location for *A. actinomycetemcomitans*, normal oral hygiene routines or oral disease state can provide easy access to the blood stream and allow hematological spread of the bacterium (19–21). The resulting transient bacteremia can allow *A. actinomycetemcomitans* to bind to exposed extracellular matrix components due to previous tissue damage and lead to extra-oral disease, e.g., infective endocarditis (19–21).

The extracellular matrix binding protein A (EmaA) is a well characterized collagen binding adhesin that has been implicated in mediating the binding of *A. actinomycetemcomitans* to the exposed matrix components of damaged heart valves and the subsequent development of infective endocarditis (22). The adhesin is composed of three identical protein subunits that form antenna-like structures associated with the surface of the bacterium (23, 24). Two molecular isoforms of the adhesin have been identified, corresponding to *A. actinomycetemcomitans* strain-specific serotype: the canonical, full-length 202 kDa isoform (principally associated with serotypes b and c) and the 173 kDa isoform (serotypes a and d) (25). In addition to collagen binding activity, this adhesin is also associated with cell-to-cell interaction in biofilm formation (26). Stability of the 3D EmaA structure on the surface of the bacterium is dependent on the expression of a functional WaaL ligase (6, 7, 27–29). Curiously, the collagen binding activity of the canonical, full-length isoform is dependent on the ligase activity, whereas the 173 kDa isoform is not (7, 27). Biofilm formation activity is universally independent of ligase activity (26).

To investigate the role of WaaL in the collagen binding activity of EmaA, we have determined that *waaL* (*rflA*) of *Escherichia coli* expressed *in trans* in an *A. actinomycetemcomitans waaL* mutant strain does not restore collagen binding activity to the strain. Based on this finding, we have used two strains of *E. coli* with differing O-PS structures, neither of which bind collagen, to develop a model system to independently study the role of *A. actinomycetemcomitans* WaaL in the collagen binding activity of EmaA. Only *E. coli* strains expressing both *A. actinomycetemcomitans waaL* and *emaA* were able to effectively bind collagen and displayed biochemical changes to the EmaA monomers. These data demonstrate that the *A. actinomycetemcomitans waaL* is required for collagen binding activity associated with EmaA and suggests that this ligase is important for conferring changes in the structure of this adhesin.

## Materials and Methods

### Bacterial strains and growth conditions

Bacterial strains and plasmids used in this study are listed in Table 1. *A. actinomycetemcomitans* strains were grown from frozen stocks on solid TSBYE medium (3.0% tryptic soy broth, 0.6% yeast extract, 1.5% agar; Beckton Dickinson, Franklin Lakes, NJ) in a humidified 10% CO_2_ atmosphere at 37°C. A single colony was used as the inoculum for all experiments and the cultures were grown statically in TSBYE broth. Plasmids were maintained in *A. actinomycetemcomitans* strains by incorporation of 1.0 μg/ml chloramphenicol in the growth medium. *Escherichia coli* strains were grown from frozen stocks in LB medium (1.0% tryptone, 0.5% yeast extract, 0.5% NaCl; Beckton Dickinson) at 37°C in ambient air with agitation. *E. coli* strains containing plasmids were maintained at the following antibiotic concentrations: 100 μg/ml ampicillin; 20 μg/ml chloramphenicol; 50 μg/ml kanamycin.

**Table 1.**
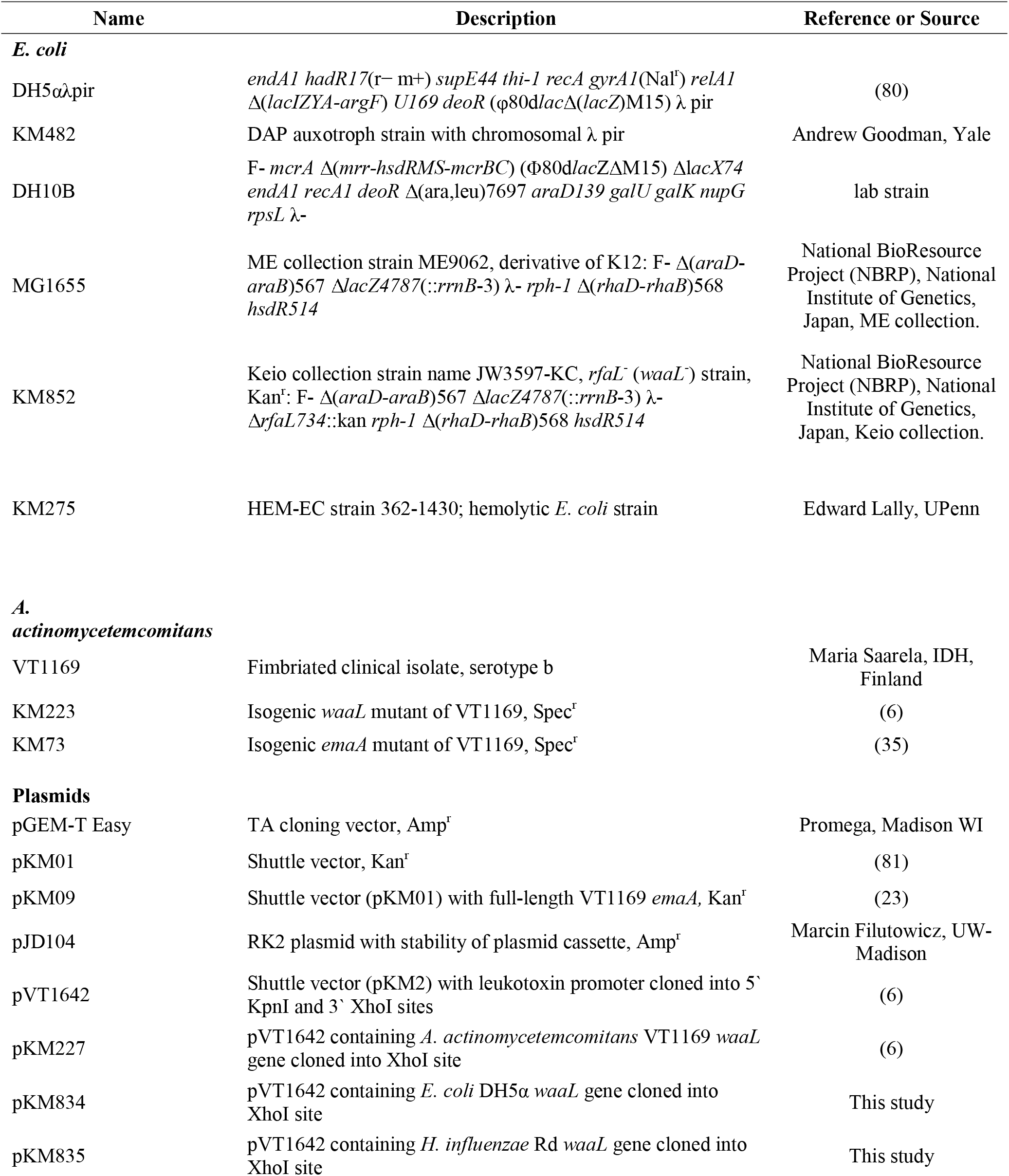
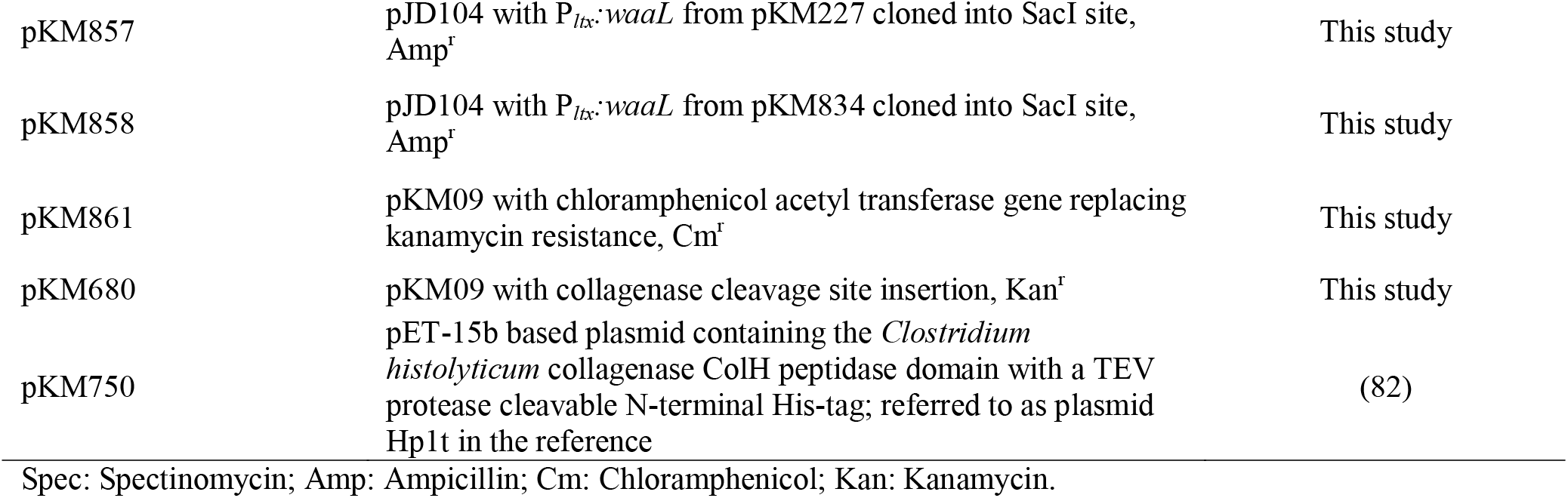
Bacterial Strain or Plasmid.

### Protein structure prediction and analysis

Tertiary structure predictions for the WaaL/RfaL proteins of *E. coli* DH5α (Accession number WP_001395405.1), *H. influenzae* Rd (Accession number AAC22538.1), and *A. actinomycetemcomitans* VT1169 (Accession number AMQ93020.1) were performed using three independent prediction algorithms, Phyre2 (30), iTASSER (31), and RaptorX (32). Structural alignments were performed by RaptorX (33, 34), and the multiple sequence alignment scores are presented in Table 3.

### Construction of *waaL* expression vectors for studies in *A. actinomycetemcomitans*

The *E. coli rfaL/waaL* gene (hereafter referred to as *waaL*) was amplified by polymerase chain reaction (PCR) using isopropanol extracted DH5α genomic DNA as the substrate and the primers ecwaaLfwdxhoI and ecwaaLrevxhoI (Table 2). The 1,260 bp amplicon was ligated into pGEM-T Easy (Promega, Madison, WI), and propagated in *E. coli* DH5α cells. The plasmid was isolated (GeneJET Miniprep Kit (ThermoScientific, Waltham, MA)), treated with with XhoI to release the amplicon and gel purified (QIAquick Gel Extraction kit (Qiagen, Hilden, Germany)). The fragment was ligated into the expression vector, pVT1642 (6), at the leukotoxin promoter 3′ XhoI site. The resulting plasmid was transformed into the *A. actinomycetemcomitans waaL* mutant strain by electroporation (35).

**Table 2.**
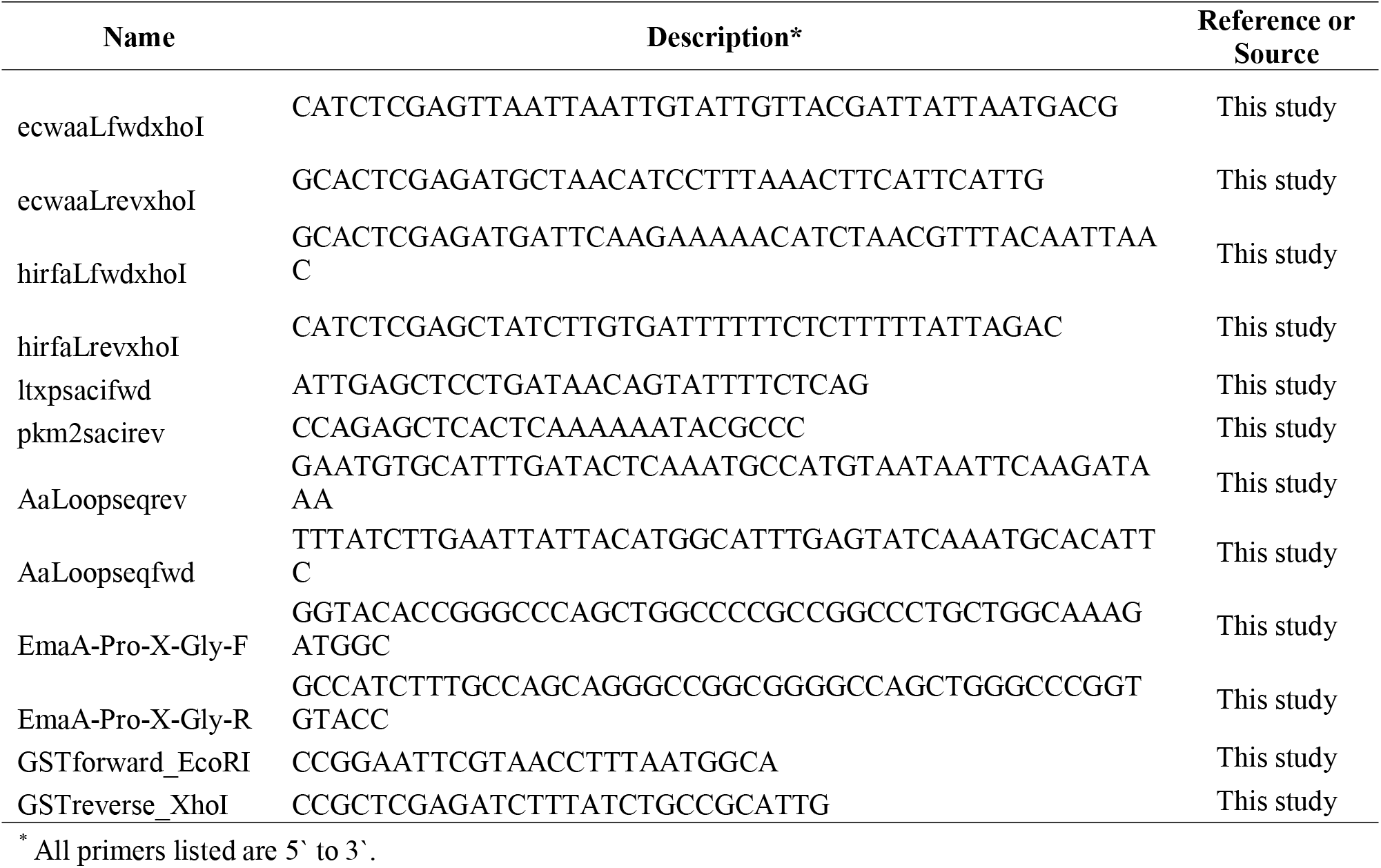
Primers.

The *H. influenzae rfaL* gene (hereafter referred to as *waaL*) was PCR amplified using isopropanol extracted genomic DNA purified from strain Rd as the substrate and the primers hirfaLfwdxhoI and hirfaLrevxhoI (Table 2). The 1,227 bp gene was cloned and transformed into *waaL* mutant *A. actinomycetemcomitans* as described above. Both the *E. coli* and *H. influenzae waaL* plasmids were confirmed by sequencing at the Advanced Genome Technologies Core facility at the University of Vermont.

The construction of the plasmid expressing the *A. actinomycetemcomitans waaL* controlled by the leukotoxin promoter has been described previously (6).

### Plasmid compatibility assay

Co-expression of both the *emaA* and the *waaL* genes in *E. coli* strains was performed using multiple plasmids. Potentially compatible combinations of plasmids with different replication systems were co-transformed into DH5α *E. coli* to check for compatibility. LB agar plates with appropriate antibiotic combinations were used to select for any colonies expressing both plasmids. pKM01, an incompatibility group B shuttle plasmid derived from pDMG4 (23), was found to be compatible with pJD104, an incompatibility group P plasmid with an RK2 derived mini replicon.

### Construction of *waaL* and *emaA* expression vectors for studies in *E. coli*

Expression of the *emaA* gene in both *E. coli* strains was accomplished using pKM09, a previously constructed kanamycin-resistant plasmid based on pKM01 (23).

The *A. actinomycetemcomitans* and *E. coli waaL* expression plasmids for transformation into the *E. coli* wild type and *waaL* mutant strains were generated by amplification of the respective leukotoxin promoter-*waaL* gene constructs described previously using the primers ltxpsacifwd and pkm2sacirev (Table 2). These amplicons were purified, digested with SacI, and ligated with pJD104 previously treated with the same enzyme. The resulting plasmids were transformed into the *E. coli* strains for investigation.

### Construction of the *E. coli*/*A. actinomycetemcomitans waaL* chimera

Replacement of the predicted beta sheet of the *E. coli waaL* enzyme (corresponding to amino acids 157 - 170) with the identical region of the *A. actinomycetemcomitans waaL* enzyme was accomplished with an overlapping primer set containing the 46 base pair modified sequence. First, the 5′ end of the *E. coli waaL* gene was amplified from the expression plasmid using the primer ecwaaLfwdxhoI and the modified overlapping primer AaLoopseqrev to create a 511 base pair amplicon. Next, the 795 base pair 3′ end of the gene was PCR amplified from the same expression plasmid using the overlapping primer AaLoopseqfwd and the primer pkm2sacirev (Table 2). These amplicons were ligated together using the Gibson Assembly Master Mix (New England Biolabs, Ipswich MA) and the construct was re-amplified with ecwaaLfwdxhoI and ecwaaLrevxhoI primers to generate the modified 1,260 base pair amplicon; the amplicon was subsequently ligated into pGEM-T Easy. After propagation in *E. coli* DH5α cells and plasmid isolation, the sequence was excised using XhoI and ligated into the expression vector pVT1642 as described above. Sequence fidelity was confirmed at the Advanced Genome Technologies Core facility at the University of Vermont. The *E. coli waaL* chimera was transformed into the *A. actinomycetemcomitans waaL* mutant strain for investigation.

### Characterization of O-PS complementation

BactoELISA assays were performed to confirm the expression of *waaL* genes from various genera in both the *A. actinomycetemcomitans* and *E. coli* strains (36). 10-fold serial dilutions of 10^8^ CFU/mL bacterial cells in TSBYE or LB, respectively, were added to a 96-well plate (Nunc, Roskilde, Denmark) and allowed too fully dry overnight. The wells were washed with Tris-buffered saline (TBS: 20 mM Tris-HCl, 150 mM NaCl, pH 7.5) prior to blocking with 0.5% bovine serum albumin (BSA)-TBS. Bound bacteria were detected using whole cell rabbit polyclonal antisera: anti-serotype b *A. actinomycetemcomitans* (37), whole cell anti-*E. coli* DH5α (Novus Biologicals, Centennial, CO [NB200-579]), or whole cell anti-antigenic *E. coli* (Novus Biologicals, Centennial, CO [NB100-62526]), at 1.0 μg/mL in 0.5% BSA-TBS. The bound antibody was detected using 1:10,000 dilution of horseradish peroxidase (HRP)-conjugated goat anti-rabbit secondary antibody (Jackson Laboratory, Bar Harbor, ME). Immune complexes were detected by the addition of 100 μL/well of peroxidase substrate (4.0 mg *o*-phenylenediamine, 4.0 μL 30% H_2_O_2_, 10 mL citrate phosphate buffer (0.1 M Na_2_HPO_4_, 0.05 M Na_3_C_6_H_5_O_7_, pH 5.0)) and the reaction was terminated by the addition of 50 μL/well of 4.0 M sulfuric acid. The absorbance was quantified at a wavelength of 490 nm using an ELx800 plate reader (BioTek, Winooski, VT). Experiments were performed in triplicate with at least three biological replicates, and a one-way ANOVA test was used to identify statistical significance (*p* < 0.05).

### Collagen binding assay

The binding of EmaA to human placenta type V collagen was tested using the method as described by Yu *et al*., with modifications (24). Briefly, 100 μl of type V collagen in 0.5 N acetic acid was diluted using sodium bicarbonate buffer (16 mM sodium carbonate, 34 mM sodium bicarbonate, pH 9.6) and added to a 96-well plate before overnight incubation at 4°C. Plates were washed three times with TBS prior to blocking the wells with 0.5% BSA-TBS. 100 μl of 10^8^ CFU/mL *A. actinomycetemcomitans* cells (diluted in TSBYE) or *E. coli* cells (diluted in phosphate buffered saline (PBS: 136.9 mM NaCl, 8.1 mM Na_2_HPO_4_, 2.68 mM KCl, 1.46 mM KH_2_PO_4_, 0.46 mM MgCl_2_, pH 7.4)) were added to the wells and incubated at 37°C for one hour. Bound bacteria were detected using the immunological methods described above. Experiments were performed in triplicate with at least three biological replicates, and a one-way ANOVA test was used to identify statistical significance (*p* < 0.05).

### *E. coli* biofilm assay

Biofilm assays were based on the method of Merritt *et al*. (38). *E. coli* strains were initially grown from frozen stocks on LB agar, and isolated colonies were inoculated into 5 ml of LB broth and grown with shaking at 37°C for 16 hours. Cultures were diluted 1:100 and grown to an OD_600nm_ = 0.3; 100 μl aliquots of a 1:1000 dilution were added to a 96-well plate and the cultures were grown for 24 hours at 37°C. After, the supernatants were aspirated and any nonadherent cells were removed by three consecutive washes with TBS. Biofilms were stained with 0.1% crystal violet in water for 20 min, washed three times with TBS, and solubilized using a 2:1 solution of water:glacial acetic acid. The relative biofilm mass of each strain was quantified by absorbance at 562 nm using the ELx800 plate reader. Experiments were performed in triplicate and a minimum of three biological replicates were assayed; a one-way ANOVA test was used to identify statistical significance (*p* < 0.05).

### Surface expression of EmaA in *E. coli*

The presentation of the EmaA adhesin on the surface of *E. coli* cells was determined by enzymatic cleavage of the protein by collagenase from *Clostridium histolyticum* followed by immunoblot analysis. Collagenase cleaves the peptide sequence (Gly--Pro--X)_n_, where X is a neutral amino acid; three consecutive GPX cut sites were introduced into the *emaA* sequence corresponding to amino acids 899-907 (G**TN**G**AN**G**TD**). This was achieved by site-directed mutagenesis (QuikChange II Site Directed Mutagenesis Kit (Agilent, Santa Clara, CA)) using the primers EmaA-Pro-X-Gly-F and EmaA-Pro-X-Gly-R (Table 2) which generated the sequence G**PA**G**PA**G**PA**; sequence fidelity was verified by sequencing at the Advanced Genome Technologies Core facility at the University of Vermont.

The integrity of this sequence for susceptibility to collagenase was assessed using a Glutathione-S-Transferase (GST) fusion protein corresponding to amino acids 827 to 1190 of the EmaA peptide sequence. A 1.1 kbp amplicon of the gene encompassing the modified sequence was amplified by PCR using the primers GSTforward_EcoRI and GSTreverse_XhoI (Table 2), digested with EcoRI and XhoI, and ligated into the pGEX-6P-1 expression vector (Millipore Sigma, Burlington, MA). The plasmid containing the fusion protein was transformed into *E. coli* BL21-DE3 cells for autoinduction and purification of the 63.4 kDa peptide (39). 300 μl of the purified peptide was incubated with 162.5 units of commercially available *C. histolyticum* collagenase mixture (Sigma) in 300 mM NaCl, 20 mM 4-(2-hydroxyethyl)-1-piperazineethanesulfonic acid (HEPES), 10 mM CaCl_2_, 50 uM ZnCl_2_ (total volume of 2.0 ml) with rotation overnight at 37°C. Proteins were separated by polyacrylamide gel electrophoresis (PAGE) under denaturing conditions and EmaA was identified by immunoblotting (25).

The commercially available collagenase was found to be insufficient for *in vitro* cleavage of EmaA from the bacterial membrane. Therefore, the peptidase domain of collagenase ColH, was purified following the method of Ducka *et al*. (40). Expressed protein was bound to NiNTA beads (ThermoScientific) and eluted with 250 mM imidazole following a pre-elution of 40 mM imidazole. The protein was dialyzed into 25 mM Tris, 50 mM NaCl, pH 7.5, made to 50% glycerol, and stored at −20°C.

Cleavage of whole EmaA was assayed in an *emaA* minus *A. actinomycetemcomitans* strain or *E. coli* strain carrying the recombinant *emaA* expression plasmid pKM680. Membrane fragments were isolated, as described previously (35), and solubilized with 1% Triton X-100. The purified enzyme was added and incubated for 16 hours at 37°C, with rocking. The reaction was stopped by the addition of 10 mM EDTA (final concentration) and the insoluble, cleaved, EmaA was collected by low-speed centrifugation. Both the pellet and supernatant were analyzed via PAGE and immunoblotting as described previously (25, 35).

### Isolation and visualization of LPS from *E. coli* strains

LPS was isolated from both *E. coli* MG1655 and hemolytic strains using a modified version of the method described in Davis 2012 (41). Briefly, 50 mL of stationary phase culture was diluted to OD_600_ = 0.5, pelleted, and resuspended in sodium dodecyl sulfate (SDS) buffer (2% β-mercaptoethanol (BME), 2% SDS and 10% glycerol in 0.05 M Tris-HCl, pH 6.8, with a small addition of bromophenol blue for color). The suspension was boiled for 15 minutes before 10 μl of a 10 mg/mL proteinase K solution was added. Following incubation overnight at 55°C, 200 μl of ice-cold Tris-saturated phenol was added and vortexed for 10 seconds. The mixture was then incubated at 65°C for 15 minutes, cooled to room temperature, and 1.0 ml of ethyl acetate was added and vortexed as before. The samples were centrifuged at 20,600 x *g* for 10 minutes and the bottom layer removed. The phenol-ethyl acetate extraction was repeated a second time, followed by a further ethyl acetate only wash step. 100 μl of SDS buffer was added to the samples, and 15 μl of each was loaded on to a 4 - 15% polyacrylamide Tris/Glycine minigel (Bio-Rad, Hercules, CA) and separated at 100 V. The LPS was visualized using the Pro-Q Emerald 300 Lipopolysaccharide Gel Stain Kit (Molecular Probes, ThermoScientific) following the manufacturer’s instructions.

### Analysis of membrane bound EmaA expressed in *E. coli* strains

The outer membrane fractions of the *E. coli* strains were prepared as described previously (35). Briefly, 500 ml of mid-log phase cells were centrifuged and resuspended on ice in 3.0 ml 10 mM HEPES, pH 7.4, with 1.0 mM phenylmethylsulfonyl fluoride (PMSF, USB Corporation, Cleveland, OH) and Pierce Protease Inhibitor with EDTA (ThermoScientific). The cells were kept ice cold and lysed using a French pressure mini cell at 9000 PSI (62,100 kPa); cell debris was removed by centrifugation at 7,650 x *g*, and the lysate resuspended in 10 mM HEPES. The resuspension was centrifuged at 100,000 x *g* for 30 minutes at 4°C, washed with HEPES, and re-centrifuged. This pellet was resuspended in ~1.5 ml of 10 mM HEPES + 0.1% (w/v) N-Lauryl sarcosine sodium salt (Sigma) and incubated at room temperature for 30 minutes. The mixture was centrifuged at 16,000 x *g* for 30 minutes, and the resulting pellet washed with 10 mM HEPES before repeating the previous centrifugation. The resulting outer membrane pellet was resuspended in 10 mM HEPES and analyzed via sodium dodecyl sulfate-polyacrylamide gel electrophoresis (SDS-PAGE) followed by Immunoblot and lectin blot.

To visualize the EmaA monomers expressed in *E. coli*, equivalent concentrations of membrane protein from each sample (as assessed via absorbance at 280nm) were prepared with a 4X Laemmeli loading buffer (0.2 M Tris-HCl, 8% (w/v) SDS, 40% (v/v) glycerol, and 0.004% (w/v) bromophenyl blue), boiled for 5 min, loaded into a 4 - 15% polyacrylamide Tris/Glycine minigel (Bio-Rad), and run at 40 V for a minimum of 16 hours at room temperature. The separated proteins were transferred to a 0.2 μm nitrocellulose membrane (Amersham Protran, Cytiva, Marlborough, MA) at 120 mA for 120 minutes at 4°C, blocked with 5% (w/v) non-fat milk, and probed with an anti-EmaA stalk monoclonal antibody (25). Immune reactive complexes were detected using HRP-conjugated goat anti-mouse IgG (Jackson Laboratory) and visualized on film (USA Scientific, Ocala, FL) using SuperSignal West Pico plus chemiluminescent substrate (ThermoFisher Scientific, Waltham, MA).

Lectin blot analysis of EmaA was performed as described in Tang 2010 (6), with modifications. Outer membrane proteins were prepared and transferred to a nitrocellulose membrane as described above. The membrane was blocked using a Carbo-Free Blocking Solution (Vector Laboratories, Newark, CA) and probed with biotinylated mannose-recognizing Concanavalin A (Vector Laboratories) for 1 hour. The membrane was washed 6 separate times in TBS-T for 5 min and then incubated with HRP-conjugated streptavidin (Vector Laboratories) for an additional hour. After a further six washes, the lectin-avidin complex was visualized as described by the EmaA immunoblot.

## Results

### Function of heterologous WaaL in *A. actinomycetemcomitans*

The unique activity of the *A. actinomycetemcomitans* O-antigen ligase *waaL* was investigated by complementation studies using two different *waaL* genes, one from a related member of the Pasteurellaceae family, *Haemophilus influenzae* serotype d strain Rd, and another from the more evolutionarily distant Enterobacteriaceae family, *Escherichia coli* strain DH5α. Amino acid sequence alignment of the *A. actinomycetemcomitans* protein with the *H. influenzae* protein revealed a high percentage of similar and conserved amino acids (62% and 41%, respectively) (Fig. 1). This was in stark contrast to alignment with the *E. coli* sequence (44% similar and 23% conserved). The predicted functional residues and motifs required for the canonical ligase activity (42–44) are present in all three sequences (Fig. 1). Two notable differences in the *E. coli* sequence from the more closely related pair are an 18 amino acid gap between residues 100 - 118 as well as a 14 amino acid gap between residues 398 - 411 (Fig. 1). The tertiary structure model of each protein was based on three separate protein structure algorithms (Phyre2 (30), iTASSER (31), and RaptorX (32)) followed by alignment with the RaptorX multiple structure alignment software (33, 34) (alignment scores are presented in Table 3). All three predicted structures were very similar, with the principal exception of a small beta sheet (amino acids 157 - 170) predicted in the *E. coli waaL* that is absent in the *H. influenzae* and *A. actinomycetemcomitans* protein structures (Fig. 2).

**Fig. 1.**
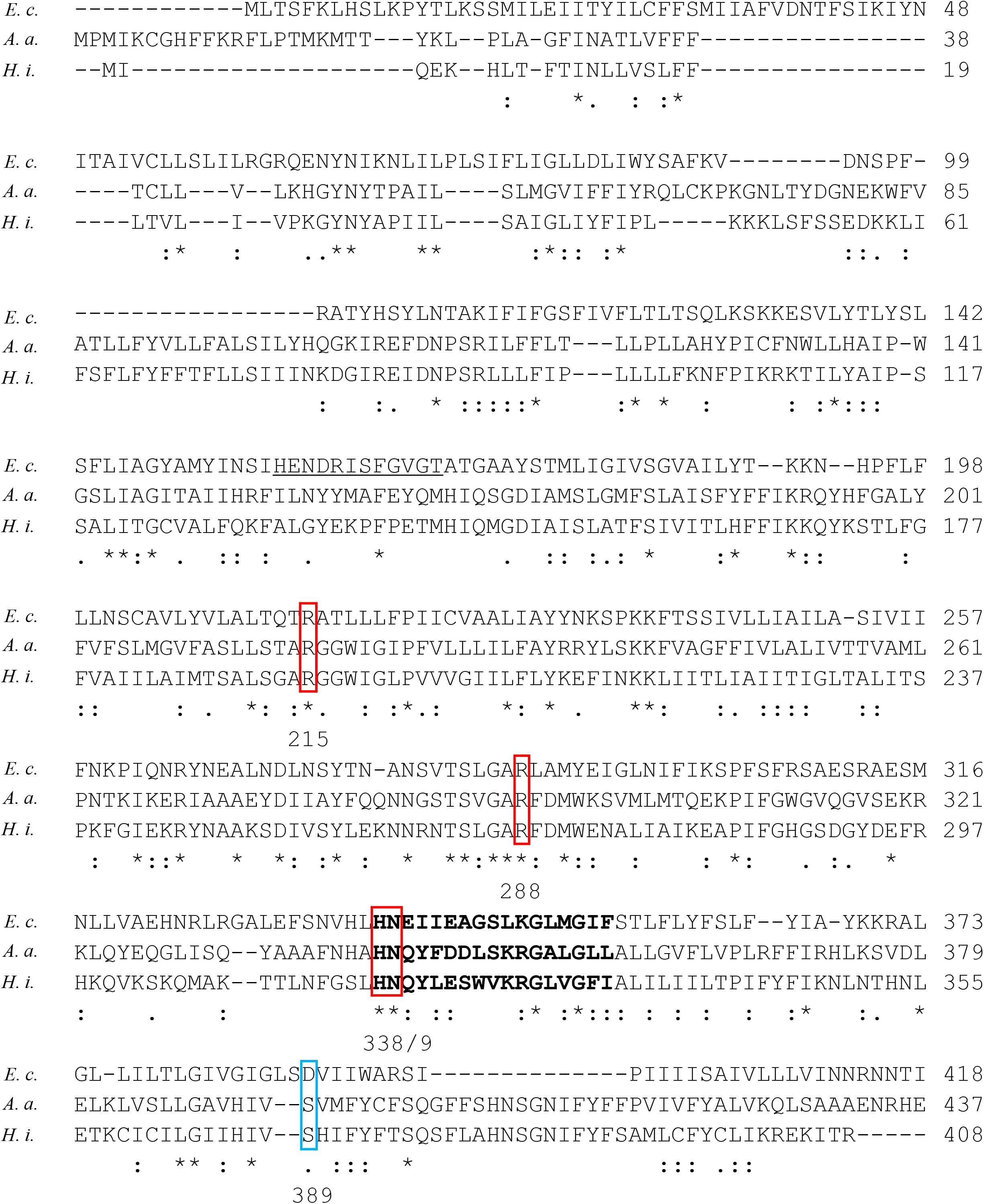
Primary amino acid sequence alignment of *waaL* orthologs from *E. coli* DH5α (*E. c*.), *A. actinomycetemcomitans* VT1169 (*A. a*.), and *H. influenzae* Rd (*H. i*.). Red boxes are proposed active site residues (215, 288, 338/339), bolded region is the H[NSQ]X9GXX[GTY] motif, underlined is the predicted *E. coli* beta-sheet domain. The blue box indicates the D389 that is suggested to be required for ligase activity in *E. coli*.

**Fig. 2.**
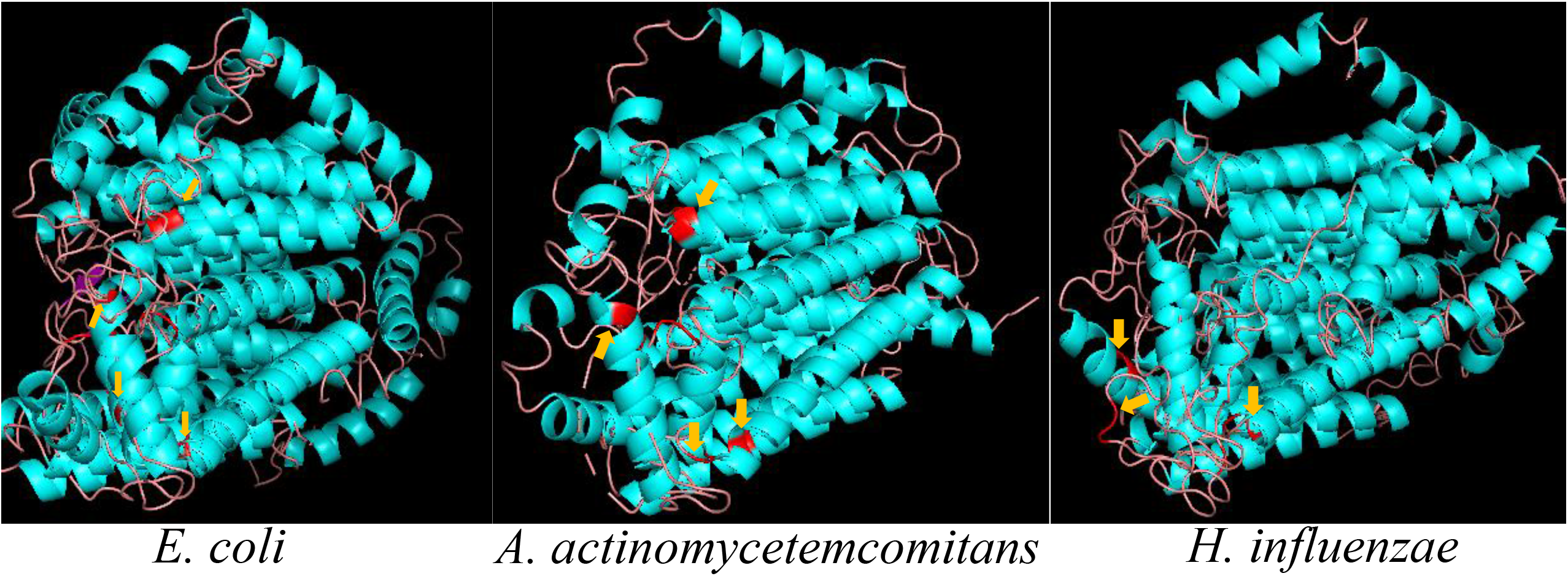
*In silico* tertiary structure predictions of WaaL orthologs of *E. coli* strain DH5α, *A. actinomycetemcomitans* strain VT1169, and *H. influenzae* strain Rd. Images consist of aligned Phyre2, iTASSER, and RaptorX models. Red (with arrows) indicates the conserved 215, 288, and 338/339 residues. Purple (circled) indicates the beta sheet found only in the *E. coli* structure.

**Table 3.** Multiple-sequence alignment scores (33, 34).

| Strain<sup>§</sup> | Lali<sup>1</sup> | RMSD<sup>2</sup> | uGDT/GDT<sup>3</sup> | TMscore<sup>4</sup> |
| --- | --- | --- | --- | --- |
| *E. c.* | 231 | 4.74 | 133/50 | 0.624 |
| *A. a.* | 253 | 4.06 | 166/59 | 0.753 |
| *H. i.* | 241 | 4.66 | 144/52 | 0.63 |

<sup>§</sup> *E. c.* (*E. coli*), *H. i.* (*H. influenzae*), *A. a.* (*A. actinomycetemcomitans*).
<sup>1</sup> *Lali*: length of core (all the fully-aligned residues).
<sup>2</sup> *RMSD*: root-mean-square deviation of the core residues.
<sup>3</sup> *uGDT/GDT*: *uGDT* = unnormalized Global Distance Test score; *GDT* = (*uGDT* / the shortest protein length) x 100.
<sup>4</sup> *TMscore*: average *TMscore* of all the pairwise alignments; >0.6 is a >90% chance that two proteins share a similar fold, <0.4 is a >90% chance of different folds.

Individual plasmids containing these O-antigen ligase genes were transformed into a *waaL* minus serotype b strain of *A. actinomycetemcomitans* (7) (which expresses the canonical full-length *emaA*) and assayed for the expression of O-PS and collagen binding activity. Transformation of the *A. actinomycetemcomitans waaL* mutant strain with the *E. coli, H. influenzae*, or native *waaL* gene all increased O-PS expression to levels equal to or greater than those observed for the parent strain (Fig. 3A). However, of the two non-native genes, only the *H. influenzae waaL* transformed strain increased the bacterial collagen binding activity to levels similar to the wild type (88.7 ± 19.0%) or the mutant strain transformed with the endogenous gene (115.7 ± 24.8%) (Fig. 3B). Transformants expressing the *E. coli waaL* gene did not have significantly different collagen binding activity from the *A. actinomycetemcomitans waaL* mutant strain (Fig. 3B). Transformation of a chimeric *E. coli waaL* (the *E. coli* sequence predicted to form a beta sheet replaced with the *A. actinomycetemcomitans* sequence, see Fig. 1 and 2) into the *A. actinomycetemcomitans waaL* mutant strain also did not restore collagen binding activity (Supplementary Fig. 1).

**Fig. 3.**
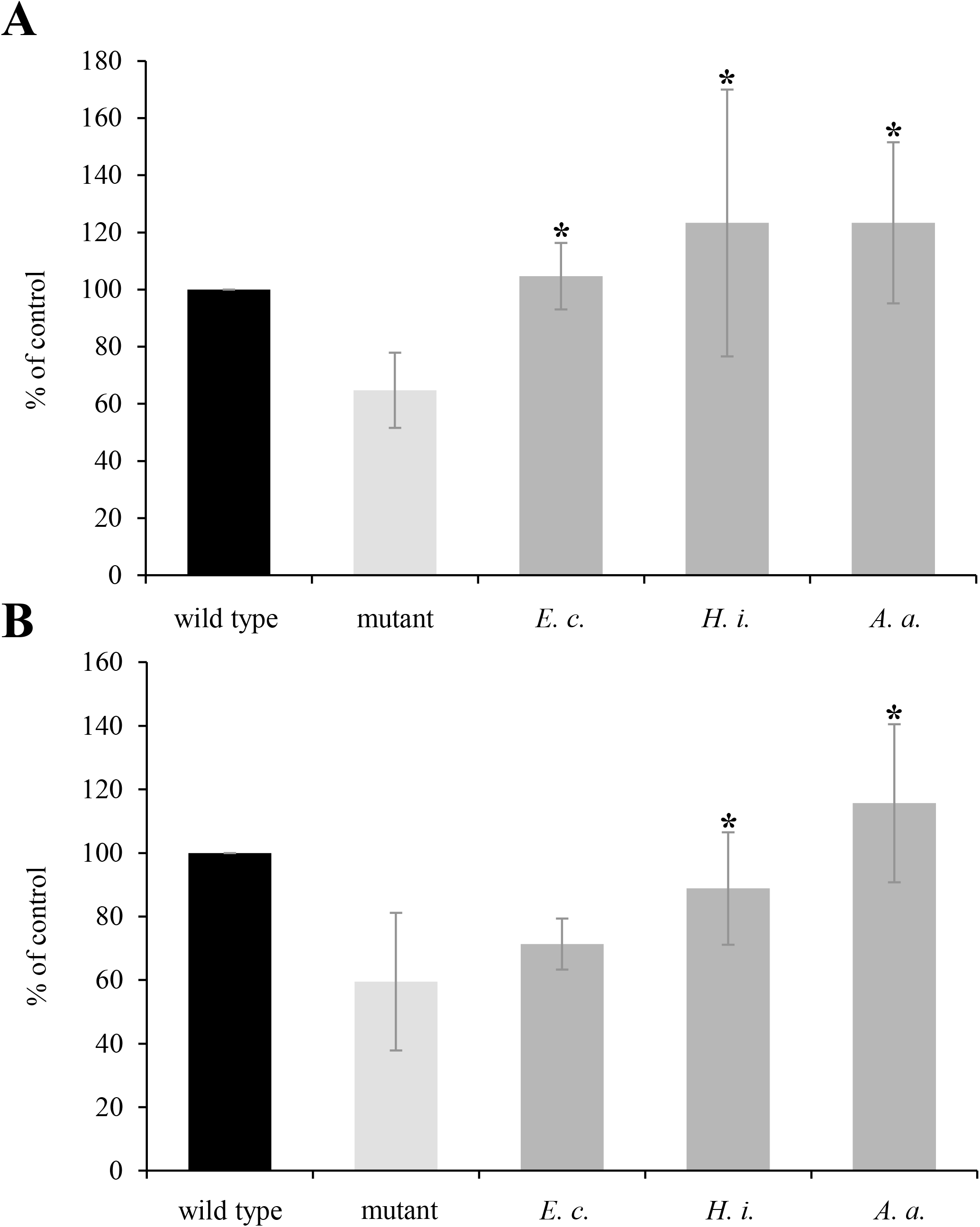
Functional activity of heterologous WaaL expressed *in trans* in *A. actinomycetemcomitans*. A) LPS biosynthesis. Strain VT1169 (wild type), VT1169 *waaL* mutant (mutant), and the mutant transformed with a plasmid expressing one of the following: *E. coli* strain DH5α *waaL* (*E. c*.), *H. influenzae* strain Rd *waaL* (*H. i*.), VT1169 *waaL* (*A. a*.). BactoELISA data was normalized to the wild type and was set at 100%; a minimum of three biological replicates were performed. B) Collagen binding activity of the same strains; data was normalized to the wild type and was set at 100%; a minimum of three biological replicates were performed. The statistical significance is indicated with an asterisk (*p* < 0.05).

### Functional expression of WaaL and EmaA in *E. coli*

The observation that the *E. coli* WaaL enzyme did not increase the collagen binding activity of EmaA allowed for the development of a model system to assess the glycosyl transferase activity of the *A. actinomycetemcomitans* WaaL with respect to the post-translational modification of EmaA. To achieve this, the collagen binding activity of EmaA was assessed by co-expression of both *emaA* and *A. actinomycetemcomitans waaL* in two *E. coli* strains: MG1655, a K12 derivative with an incomplete O-PS (45), and a hemolytic isolate (HEC) with a complete O-PS. Neither strain normally binds to collagen. Simultaneous expression of both *emaA* and *waaL* was achieved using two independent plasmids with unique, compatible origins of replication; co-expression of these plasmids did not lead to any observable growth defects in either of two *E. coli* strains. The use of dual compatible plasmids enabled phenotypic examination of each expressed protein on an individual basis.

The functionality of the *A. actinomycetemcomitans waaL* in *E. coli* was determined by transformation of a plasmid containing the nucleic acid sequence of the gene into an *E. coli* MG1655 *waaL* mutant strain (46); this strain was assayed for expression of O-PS using anti-*E. coli* DH5α antisera in an ELISA format. The immunoreactivity of the strain transformed with the *A. actinomycetemcomitans waaL* was comparable to the parent *E. coli* MG1655 strain or the *waaL* minus *E. coli* strain complemented with the *E. coli* gene (Fig. 4A). Isolation of the LPS from these strains demonstrated no observable difference in the LPS banding pattern following glycan staining (Fig. 5A).

**Fig. 4.**
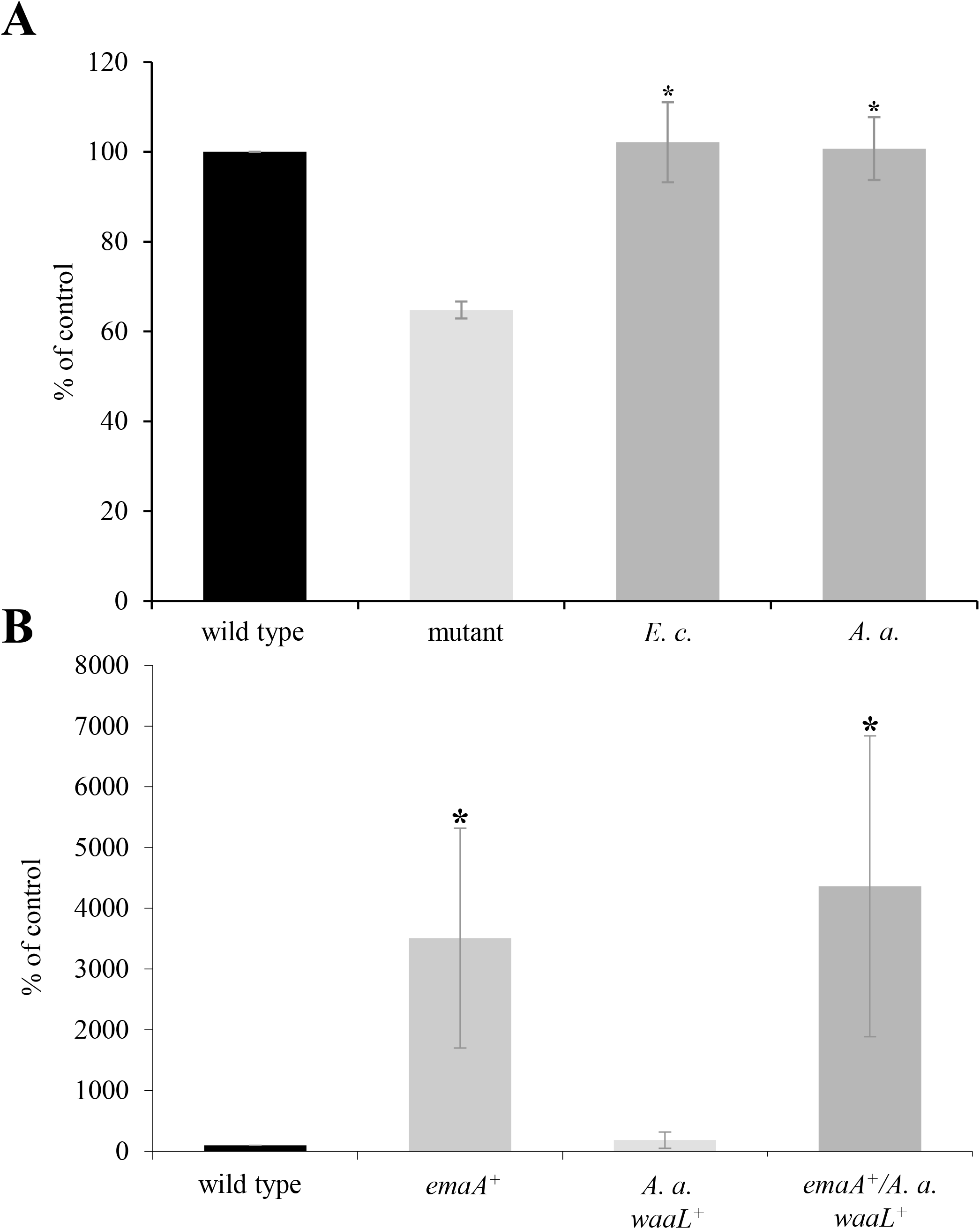
Functional activity of heterologous WaaL expressed *in trans* in *E. coli*. A) LPS biosynthesis. *E. coli* strain MG1655 (wild type), isogenic *E. coli waaL* mutant (mutant), *waaL* mutant transformed with a plasmid expressing *E. coli waaL* (*E. c*.), *waaL* mutant transformed with a plasmid expressing *A. actinomycetemcomitans waaL* (*A. a*.). BactoELISA data was normalized to the wild type and was set at 100%; a minimum of three biological replicates were performed. B) Biofilm formation activity. Wild type (MG1655), *emaA^+^* (transformation with plasmid expressing *emaA*), *A. a. waaL^+^* (transformation with plasmid expressing *A. a. waaL*), *emaA^+^*/ *A. a. waaL^+^* (co-transformation with plasmids expressing *emaA* and *waaL*). Data was normalized to the wild type and was set at 100%; a minimum of three biological replicates were performed. The statistical significance is indicated with an asterisk (*p* < 0.05).

**Fig. 5.**
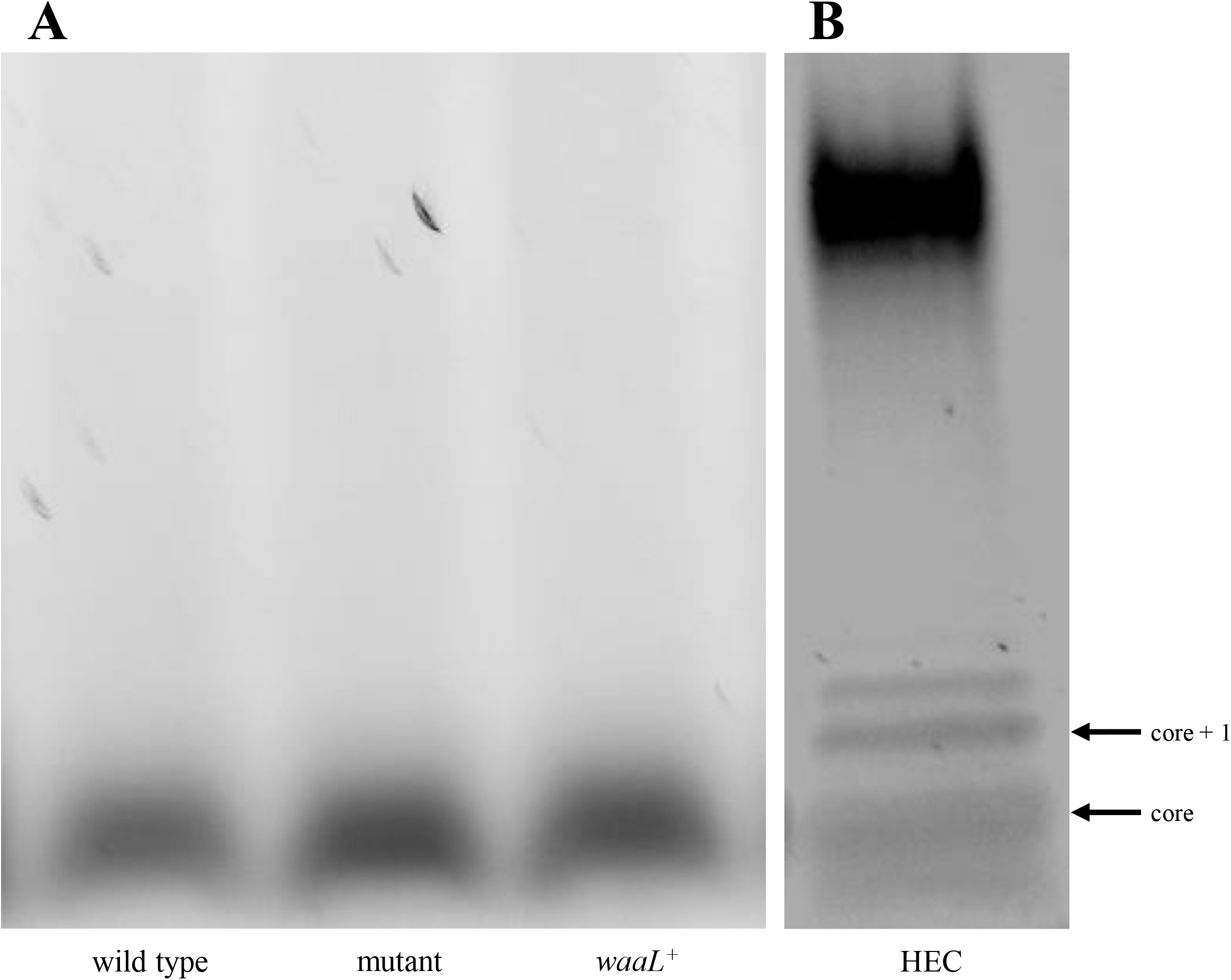
O-PS expression of *E. coli* strains. Isolated LPS from *E. coli* strains was stained with Pro-Q Emerald 300 LPS staining kit. A) Left: *E. coli* strain MG1655 (wild type), center: isogenic MG1655 *waaL* mutant (mutant), right: *waaL* mutant transformed with a plasmid expressing *A. actinomycetemcomitans waaL* (*waaL^+^*). B) Hemolytic *E. coli* (HEC). LPS core and core + 1 O-antigen repeat are indicated with arrows.

The synthesis of EmaA by the MG1655 *E. coli* strain transformed with the *emaA* expression plasmid was initially confirmed by immuno-dot blot analysis (Supplementary Fig. 2). Immunoreactive material was found to be present in the cytosolic, inner membrane, and outer membrane fractions (Fig. 6A). Localization of EmaA to the outer surface of the bacterium when expressed in *E. coli* was confirmed by proteolysis of this trimeric protein containing an engineered collagenase cleavage site and the addition of collagenase. The cognate collagenase cleavage sequence, (Gly—Pro—X)_n_, was engineered in triplicate into the *emaA* sequence corresponding to the stalk region (amino acids 899 to 907) by site directed mutagenesis. The fidelity of the cleavage site was investigated using a monomeric 63.4 kDa GST-EmaA stalk fusion protein; treatment of this fusion protein with collagenase resulted in the loss of the ~63 kDa immunoreactive protein and the appearance of the predicted 37.4 kDa species (Fig. 6B). The proper folding of the modified EmaA trimer was determined via transmission electron microscopy (TEM) (47) to form the antennae-like structures characteristic of the adhesin (23), and collagen binding activity was confirmed via expression in an *A. actinomycetemcomitans emaA* mutant strain (35) (Supplementary Fig. 3). The accessibility of the modified EmaA protein to collagenase was confirmed by incubation of *A. actinomycetemcomitans* cells expressing the modified protein with purified ColH and immunoblotting with a monoclonal antibody specific to the amino terminal portion of the protein (25). This generated the expected 87 kDa immunoreactive species based upon the known sequence (Fig. 6C). In the *E. coli* strain expressing the modified EmaA sequence, treatment with ColH resulted in a decrease in the immunoreactivity of EmaA associated with the membrane compared to the untreated cells (Fig. 6D), indicating that this protein is correctly oriented to the extracellular milieu.

**Fig. 6.**
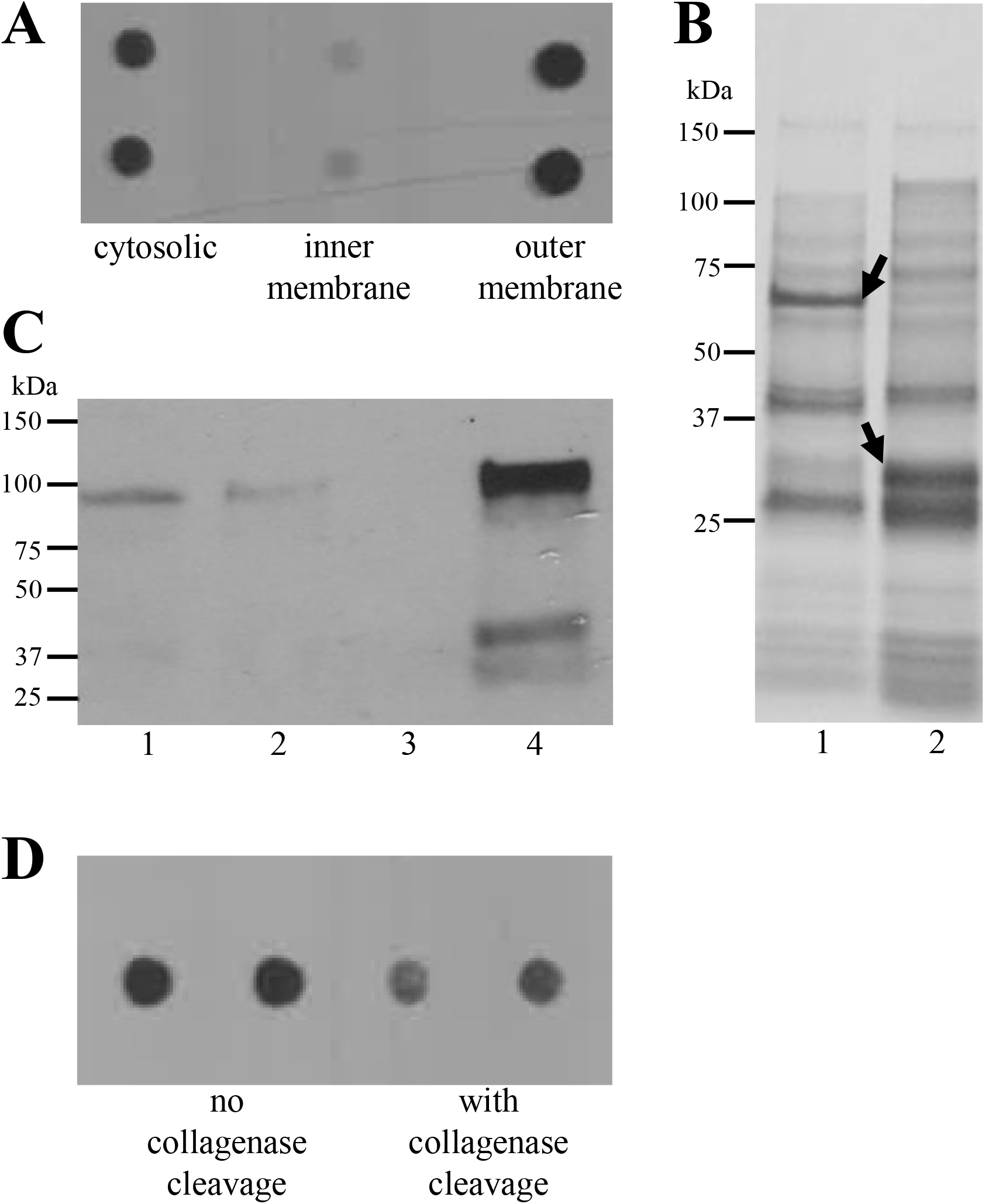
Localization of EmaA expression. A) Immuno-dot blot analysis of EmaA expression *in trans* in *E. coli*. Bacteria were fractionated and the cytosolic, inner membrane, and outer membrane fractions were investigated for EmaA using a monoclonal antibody specific for EmaA. B) Collagenase cleavage of GST-EmaA fusion protein. Fusion proteins were incubated with commercial collagenase, the products separated by PAGE, and stained with Colloidal blue. Lane 1: no enzyme, lane 2: plus enzyme. Black arrows indicate either the intact (63.4 kDa) or cleaved (37.4 kDa) protein. C) Immunoblot of the ColH cleavage product in *A. actinomycetemcomitans*. Membrane fragments were isolated from a strain expressing modified EmaA containing an engineered collagenase cleavage sequence and treated with purified ColH. The cleaved product was collected by centrifugation and both the pellet and supernatant were analyzed for immunoreactive material using a monoclonal antibody specific for the N-terminal sequence of EmaA (25). Lane 1: pellet, lane 2: supernatant, lane 3: negative control (*emaA^−^*), lane 4: supernatant fraction after concentration. D) Immuno-dot blot analysis of intact and treated membrane fraction of *E. coli* expressing modified EmaA *in trans*. Membrane fractions were isolated and incubated with or without the purified ColH.

The functionality of EmaA expressed in *E. coli* was determined by the ability of these cells to increase the mass of biofilm formed compared to the control cells, since the EmaA adhesin contributes to biofilm formation in *A. actinomycetemcomitans* independent of Waal ligase activity (26, 48). *E. coli* strain MG1655 formed very little biofilm, whereas transformation of this strain with the plasmid expressing *emaA*, under the regulation of the endogenous promoter, resulted in a greater than a 40-fold increase of the mass of the formed biofilm compared with the control strain (Fig. 4B). These results suggest that the EmaA adhesin maintains the correct quaternary structure for biofilm activity.

### Modification of EmaA monomers expressed in *E. coli*

In *A. actinomycetemcomitans*, collagen binding activity of the canonical 202 kDa isoform is dependent upon WaaL ligase and is suggested to be the enzyme required for glycosylation (6, 7). To determine if the *A. actinomycetemcomitans* WaaL ligase independently modified EmaA and increased collagen binding activity in an *E. coli* background, a modified collagen binding assay was developed using a whole cell *E. coli* polyclonal antisera, generated against a strain with the same O-PS composition as MG1655. Transformation of wild type *E. coli* MG1655 (expressing chromosomal *E. coli waaL*) with the plasmid expressing the *A. actinomycetemcomitans waaL* did not result in a statistically significant change in collagen binding activity when compared with the control strain. Minor changes in collagen binding activity were observed for the same strain transformed with the *emaA* plasmid alone. However, *E. coli* cells co-expressing both *emaA* and *A. actinomycetemcomitans waaL* increased collagen binding by over six times that of the wild type strain (Fig. 7). Similar results were observed when the plasmids expressing *emaA* and *A. actinomycetemcomitans waaL* were transformed into a hemolytic *E. coli* isolate (HEC) of unknown serotype that nonetheless synthesizes a robust O-PS (Fig. 5B).

**Fig. 7.**
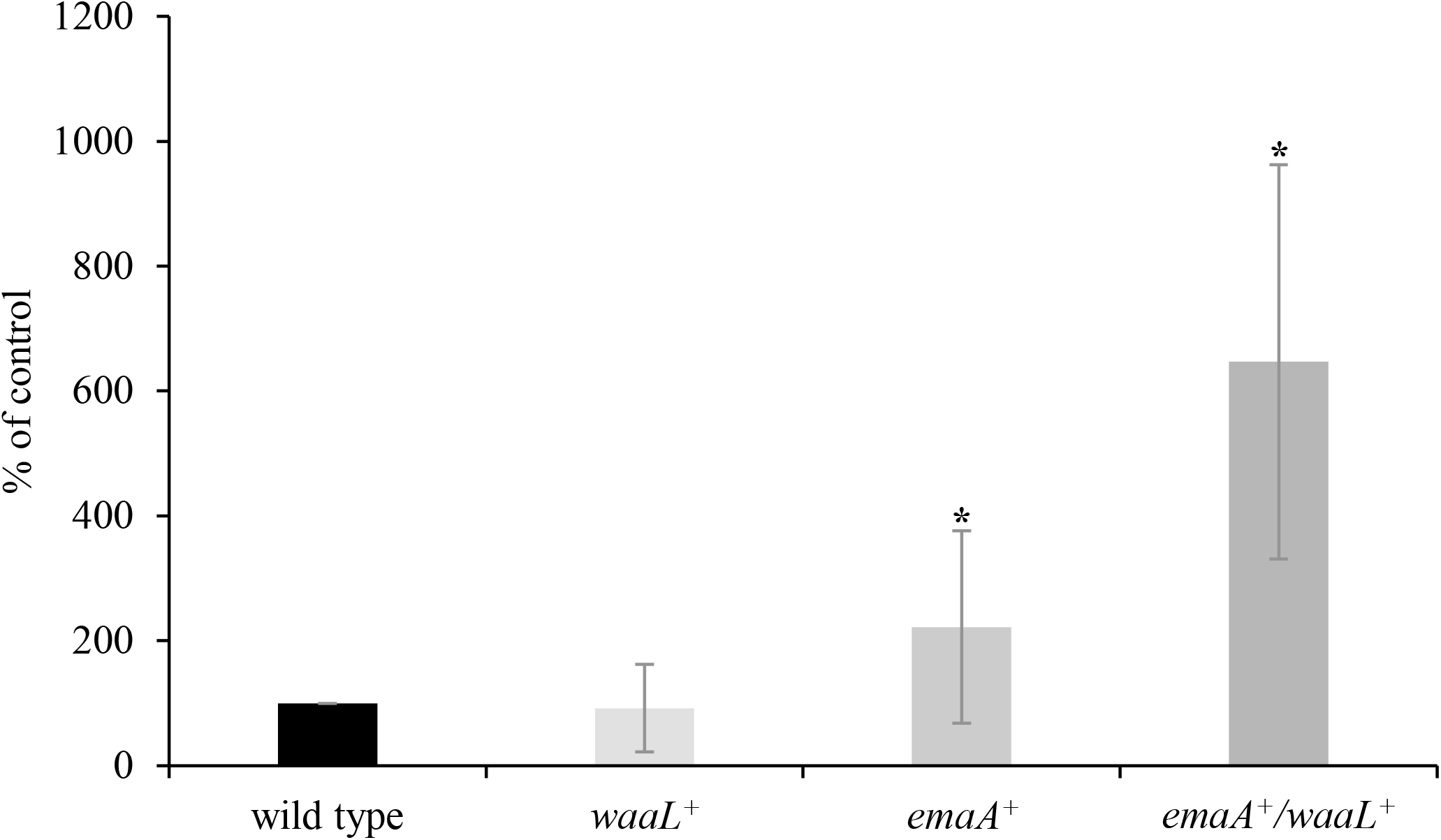
Collagen binding activity of *E. coli* strain MG1655 with *in trans* expression of *A. actinomycetemcomitans* proteins. Strain MG1655 (wild type), MG1655 transformed with a plasmid expressing *emaA* (*emaA^+^*), MG1655 transformed with a plasmid expressing *A. actinomycetemcomitans waaL* (*A. a. waaL^+^*), MG1655 co-transformed with plasmids expressing *emaA* and *A. actinomycetemcomitans waaL* (*emaA^+^*/*waaL^+^*). Collagen binding was normalized to the wild type and was set at 100%; a minimum of three biological replicates were performed. The statistical significance is indicated with an asterisk (*p* < 0.05).

The increase in collagen binding of the *E. coli* strains co-transformed with both plasmids suggests that the adhesin is modified, as observed in *A. actinomycetemcomitans*. Since purification of this adhesin has not been achieved, changes in the electrophoretic mobility of the protein and O-PS specific lectin blotting were used as evidence for biochemical changes. Immunoblot analysis of equal concentrations of outer membrane proteins isolated from the HEC parent strain, and the HEC strain transformed with the plasmid expressing *emaA*, and the HEC strain transformed with both plasmids expressing *emaA* and *A. actinomycetemcomitans waaL* demonstrated an apparent shift in the molecular mass of the EmaA monomer co-expressed with *waaL* when compared with the strain expressing *emaA* alone (Fig. 8A). In addition, electrophoretic mobility differences were also observed in higher molecular weight bands corresponding to multimers of the EmaA (Fig. 8A). As with EmaA immunoblots in *A. actinomycetemcomitans* (6, 7), the majority of the adhesin remains in the stacking gel as protein aggregates. Little immunoreactive material was observed in the lane corresponding to the HEC parent strain not transformed with a plasmid expressing *emaA*.

**Fig. 8.**
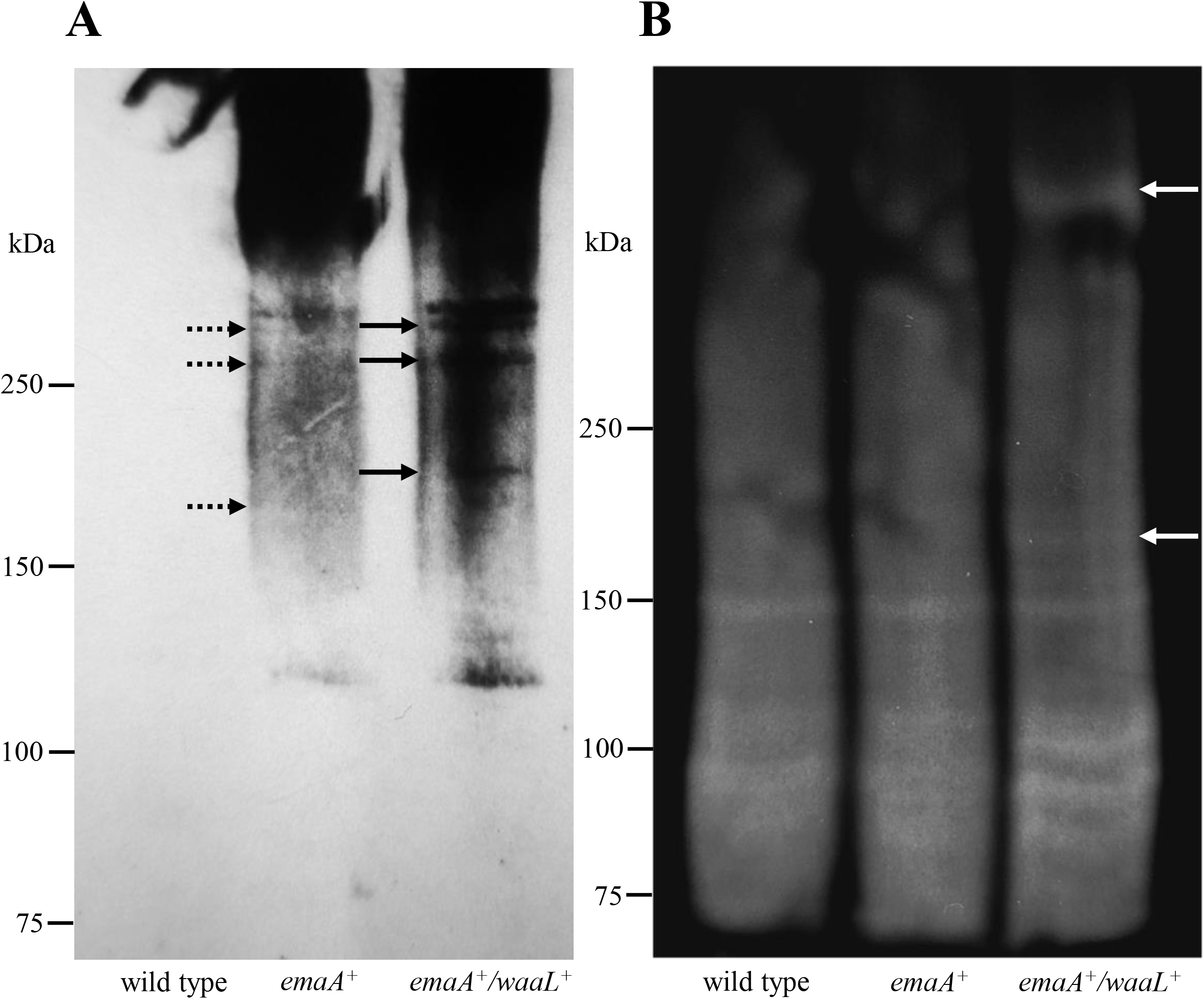
Electrophoretic mobility and glycosylation of EmaA expressed in hemolytic *E. coli* (HEC) strains. A) Immunoblot. Wild type HEC (wild type), HEC transformed with plasmid expressing *emaA* (*emaA^+^*, dashed arrows), HEC co-transformed with plasmids expressing *emaA* and *A. actinomycetemcomitans waaL* (*emaA^+^*/*waaL^+^*, solid arrows). Equivalent amounts of outer membrane fraction were isolated and separated by electrophoresis using 4–15% gradient polyacrylamide-SDS Tris-Glycine gels. The proteins were transferred to nitrocellulose and probed with an anti-stalk EmaA monoclonal antibody. The arrows indicate the electrophoretic mobility of the expressed EmaA. Note the other well defined higher molecular weight bands that also display an electrophoretic mobility shift when co-expressed with *A. actinomycetemcomitans waaL*. Immunoreactive material at the top of the immunoblot corresponds to EmaA aggregates associated with the stacking gel. B). Lectin blot. The same outer membrane fractions were prepared as above and probed with biotinylated Concanavalin A to detect glycans associated with any protein bands. Image color was inverted to make the bands easier to identify. The band corresponding to the EmaA monomers co-expressed with *A. actinomycetemcomitans waaL* are indicated by a white arrow. Note the aggregated protein near the top of the gel, also indicated with an arrow.

The presence of *E. coli* sugars associated with EmaA was detected using Concanavalin A, a lectin that recognizes the hemolytic strain of *E. coli*. As the serotype of this strain is unknown, a lectin panel was initially used to screen for lectins that would recognize the constituent O-PS. This panel included Concanavalin A, Glycine max (soybean) agglutinin (SBA), *Triticum vulgaris* (wheat germ) agglutinin (WGA), *Dolichos biflorus* agglutinin (DBA), *Ulex europaeus* agglutinin (UEA-I), *Ricinus communis* agglutinin (RCA_120_), and *Arachis hypogaea* (peanut) agglutinin (PNA); Concanavalin A was the only lectin that recognized the HEC O-PS. Using Con A, multiple staining bands were present in all three strains used in this experiment (Fig. 8B). However, an additional band was observed only in the strain co-expressing both plasmids in the region of the gel corresponding to the EmaA monomer. This band is not present in the strain expressing only chromosomal *E. coli waaL* (Fig. 8B). In addition, there was an increase in the amount of lectin staining in the region of the gel corresponding to EmaA aggregates. This staining was absent in the strain expressing only *E. coli waaL*.

The electrophoretic mobility of EmaA expressed in *E. coli* MG1655 plus or minus *A. actinomycetemcomitans waaL* was also examined. In this strain, we also observed a difference in the electrophoretic mobility of the EmaA monomer and higher molecular weight multimers (Fig. 9). The mobility difference is less than that observed in protein isolated from the HEC strain, which may be due to this strain synthesizing an incomplete O-PS. Taken together, these results demonstrate that the *A. actinomycetemcomitans* WaaL ligase is capable of changing the biochemical properties of EmaA that is required for the collagen binding ability of this adhesin.

**Fig. 9.**
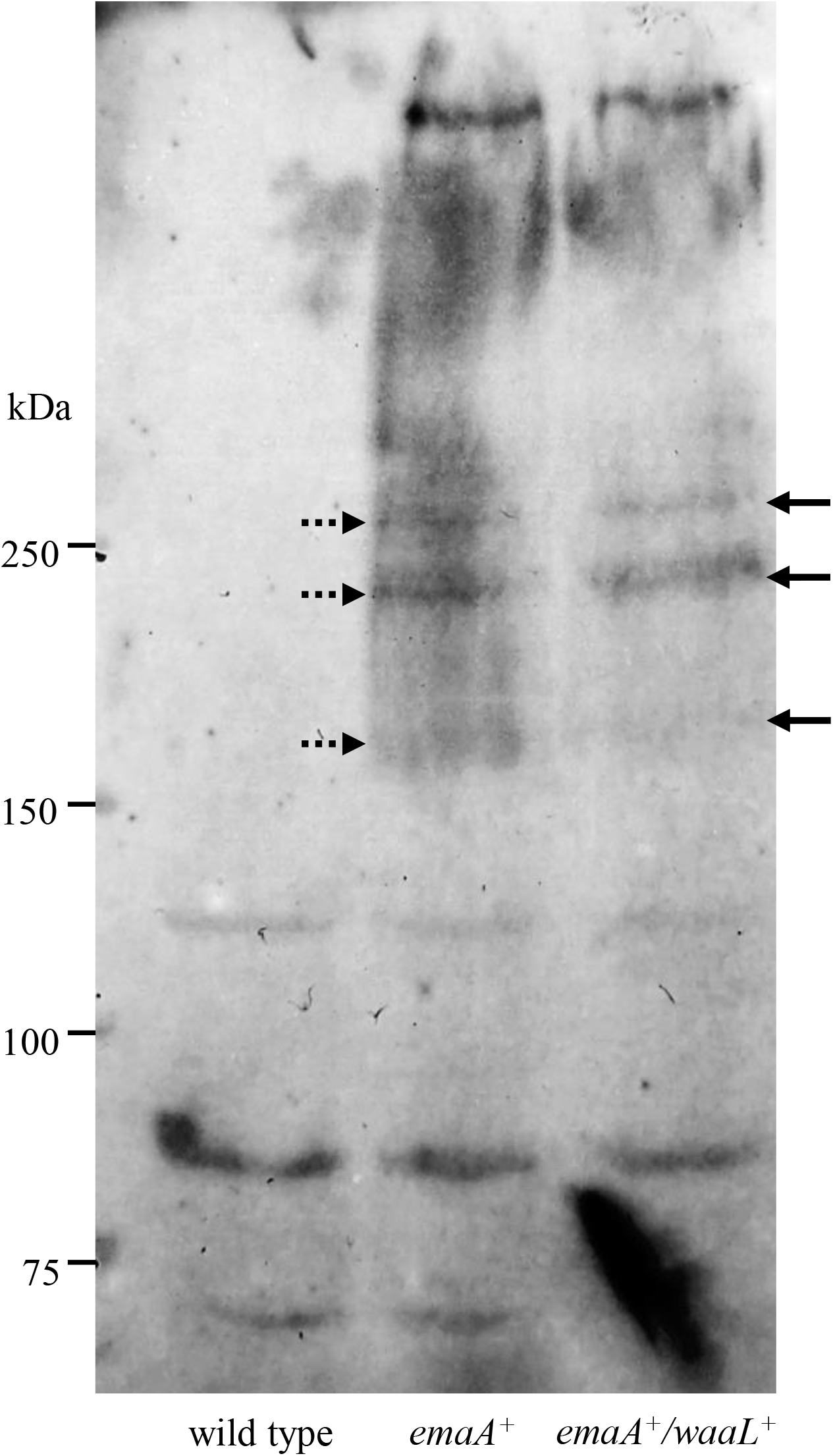
Electrophoretic mobility and glycosylation of EmaA expressed in *E. coli* MG1655 strains. Immunoblot prepared as in Fig. 8A. Strain MG1655 (wild type), MG1655 transformed with plasmid expressing *emaA* (*emaA^+^*, dashed arrows), MG1655 co-transformed with plasmids expressing *emaA* and *waaL* (*emaA^+^*/*waaL^+^*, solid arrows). The arrows indicate the electrophoretic mobility of the expressed EmaA. As with the hemolytic strain, take note of the higher molecular weight aggregates that also display an electrophoretic mobility shift when co-expressed with *A. actinomycetemcomitans waaL*.

## Discussion

The canonical function of WaaL ligase is to act as an inner membrane glycosyltransferase to mediate the transfer of the O-antigen polysaccharide (O-PS) from its undecaprenyl-diphosphate (Und-PP) linked intermediate to the core oligosaccharide of LPS (5, 42, 49, 50). Noticeable differences were apparent in the primary amino acid sequence between the two families of bacteria, whereas more similarity in sequence was observed between members of the same family. Independent of sequence differences, the predicted overall structure of these ligases is conserved and is characterized by multiple transmembrane helices and a variably sized periplasmic loop close to the C-terminus (Fig. 2) forming the catalytic center (42, 44). The periplasmic loop contains critical charged amino acids necessary for substrate binding and the ligase activity (42, 43). Four amino acids (residues R215, R288, H338, and D389 (numeration is based on the *E. coli* K12 enzyme)) and the motif H[NSQ]X9GXX[GTY] beginning with H338 have been identified to be critical for activity in most bacterial species (42–44, 51, 52). The *A. actinomycetecomitans* and *H. influenzae* sequences include three of these conserved residues as well as the motif, but do not contain the aspartic acid residue at 389 (Fig. 1). In addition, a 14-residue divergence in the sequence adjacent to this site is not present in the *E. coli* sequence, nor is a similarly sized divergence (18-residue) near the amino terminus. The *E. coli* sequence contains a predicted beta-sheet domain corresponding to amino acids 157 to 170 that is absent from the other two bacteria.

The observation that all three enzymes restored O-PS synthesis when expressed in *A. actinomycetecomitans* (Fig. 3A) suggests that these and other sequence differences do not directly impact the O-PS ligase activity associated with the different WaaL proteins. However, the inability of the *E. coli* O-antigen ligase to restore collagen binding activity in a *waaL* mutant strain of *A. actinomycetemcomitans* (Fig. 3B) suggested that the *E. coli* enzyme does not recognize EmaA as an acceptor molecule in the glycosyltransferase reaction. Some of the sequence differences described above may account for the ability of these ligases to recognize EmaA as the acceptor molecule for the Und-PP glycans. The proximity of the beta sheet domain in the *E. coli waaL* enzyme to the predicted catalytic center (Fig. 2) suggested that this sequence may be important for the recognition of EmaA as an acceptor molecule for glycosyltransferase activity. However, an *E. coli* chimera enzyme containing the *A. actinomycetemcomitans* loop at this position lacked the ability to restore collagen binding in the *A. actinomycetemcomitans waaL* mutant. It is notable that the evolutionarily more closely related enzyme from *H. influenzae* also increased collagen binding activity, indicating that the noncanonical WaaL protein activity is not exclusive to the *A. actinomycetemcomitans* ligase, but may be shared with other members of this family of bacteria.

The acceptor core oligosaccharide of the organisms used in this study vary, and yet all three individual ligases complement the O-PS defect in *waaL* mutant strains. This suggests that the WaaL homologs lacked specificity for the core acceptor oligosaccharide. A similar laxity in acceptor molecule preference has previously been observed for *E. coli* K12 WaaL (53); the *E. coli waaL* utilized in this study was obtained from *E. coli* strain DH5α, a derivitave of K12 (45). Additionally, this finding demonstrates that the WaaL homologs used in this study lack specificity for the Und-PP glycan donor substrate, supporting observations seen of other O-antigen ligases (5, 49, 54, 55), and suggests that the structure of the glycan is not important for recognition by WaaL. The *E. coli* MG1655 O-PS is a straight chain of four repeated sugars (D-galactose, D-glucose, L-rhamnose, and D-N-acetylglucosamine (56)) while the *A. actinomycetemcomitans* O-PS is a branched chain of three repeating sugars (D-fucose, L-rhamnose, and D-N-acetylgalactosamine (57)). Although the MG1655 strain of *E. coli* does not synthesize a complete endogenous O-PS due to inactivation of rhamnose biosynthetic pathway (58), the increased immunoreactivity of the transformed strain (Fig. 4A) suggested that some Und-PP-linked sugars are available for ligation, possibly non-polymerized dimers of D-galactose and D-glucose. Both the *E. coli* and the *A. actinomycetemcomitans* O-antigen ligases have the capacity to recognize the divergent Und-PP glycans when expressed in the opposing strain background. Furthermore, our recent studies suggest that the mere presence of a glycan, not the specific type, is the paramount factor for maintaining EmaA structure and function (27, 59). This observation, coupled with the lack of the *E. coli* enzyme recognition for EmaA as an acceptor and the fact that the *A. actinomycetemcomitans* enzyme interacts with the available *E. coli* sugars, make *E. coli* a suitable model system to directly investigate this novel dual function of the *A. actinomycetemcomitans* WaaL.

We reconstituted the EmaA modification system into *E. coli* cells by co-expressing the *A. actinomycetemcomitans emaA* and *waaL* genes *in trans* on independently replicating plasmids. The canonical function of *A. actinomycetemcomitans* WaaL to ligate O-PS to the core polysaccharide of LPS was confirmed by increased immunoreactivity of the transformed strain when compared with the *E. coli waaL* mutant, to a level similar to the parent strain (Fig. 4A), even if this was not accompanied by an observable change in the O-PS profile (Fig. 5A). Expression of *emaA* in another *E. coli* strain has been shown to enhance the biofilm potential of the cell (48). In both of the *E. coli* strains used in this study (MG1655 and a hemolytic strain synthesizing a complete O-PS) we also observed an increase in biofilm generated when EmaA was expressed (Fig. 4B), which suggested that the adhesin was properly expressed and localized to the outer leaflet of the membrane. This was confirmed by immuno-dot blots of cellular fractions, as well as by the decrease in immunoreactive staining of membrane bound EmaA following treatment of the intact bacterium with collagenase (Fig. 6D). Together, the data indicated that the *A. actinomycetemcomitans* genes are properly expressed and function in a similar manner when expressed in *E. coli*.

The ability of the canonical, full-length isoform of EmaA to mediate collagen binding in *A. actinomycetemcomitans* is dependent upon the synthesis of O-PS components and the presence of the O-antigen ligase WaaL (6). Therefore, *E. coli* transformants expressing this isoform of EmaA on their surface were predicted to display an increased ability to bind collagen only when co-expressed with *waaL* from *A. actinomycetemcomitans*. Although we did not observe any statistically significant increase in collagen binding of transformants expressing only *A. actinomycetemcomitans waaL*, a minor but significant increase was observed for transformants expressing only *emaA*. We attribute this increase to the auto-aggregation phenotype due to cell-to-cell interaction between EmaA expressing cells (26). The presentation of EmaA on the cell surface can generate aggregates of cells resulting in an increased signal in the assay compared to the control and does not represent individual cells binding directly to collagen. This increase in apparent collagen binding, however, was dwarfed by the collagen binding activity of cells transformed with both expression plasmids (Fig. 7) and was observable in both *E. coli* strain backgrounds. As all *E. coli* strains used for this assay were expressing chromosomal *E. coli waaL*, this gain-in-function of collagen binding by the bacteria can therefore be attributed to EmaA monomers acting as acceptor molecules in the *A. actinomycetemcomitans* WaaL glycosyltransferase reaction, using the *E. coli* Und-PP-glycans as the substrate. In addition, the difference in the O-PS composition of *E. coli* (56) and *A. actinomycetemcomitans* (57) confirms that the collagen binding activity of the canonical full-length EmaA isoform is independent of the specific composition of the added oligosaccharide(s), but is strictly dependent on the 3D conformation of the adhesin as induced by the presence of glycans (7, 29).

The biochemical analysis of the identity of the glycans associated with EmaA has been stymied by the aggregation phenotype of this adhesin, confounding our attempts for direct evidence for EmaA glycosylation. Recovery of full-length EmaA via cell lysis yields frustratingly small amounts of monomer, such that serial collection and concentration is impractical for biochemical analyses. Collagenase treatment of cells expressing a collagenase cleavage site engineered into the *emaA* sequence results in a product that can be recovered by low-speed centrifugation, suggestive of a cleavage product that is most likely forming large, high molecular weight aggregates. These aggregates can only be partial solubilized in high concentrations of SDS or 6M guanidine hydrochloride (data not shown). These aggregates likely explain the presence of immunoreactive material that does not enter the separating gel and the limited amount of monomer that is detected in the immunoblots.

In *A. actinomycetemcomitans* strains, the monomers of both isoforms of EmaA (the canonical 202 kDa and the truncated 173 kDa) display an electrophoretic mobility shift when the O-PS biosynthetic pathways of the strains are disrupted (6, 7, 27). Furthermore, O-PS specific lectin blots suggest that EmaA is associated with glycans similar to the O-PS of the expressing *A. actinomycetemcomitans* strain (6). In the absence of a functional WaaL, there are changes in the 3D structure of the functional domain of the canonical EmaA adhesin (28, 29) and a corresponding decrease in the amount of the adhesin located on the outer membrane (7). More recently, we have observed subtle changes in the structure, but not the function, of this isoform when expressed in a noncognate serotype (59). In this study, we demonstrated that *emaA* can be functionally expressed in *E. coli* only when co-expressed with *A. actinomycetemcomitans waaL*, as the *E. coli* enzyme does not have the activity necessary to restore the collagen binding phenotype classically associated with this adhesin. Changes in the electrophoretic mobility of the monomers and multimers of EmaA expressed by *E. coli* are only observed when the strain co-expresses *A. actinomycetemcomitans waaL*. Furthermore, *E. coli* O-PS specific lectin reactive species are only observed when *emaA* and *A. actinomycetemcomitans waaL* are co-expressed. These results strongly suggest a relationship between ligase activity and biochemical changes of the protein that are required for collagen binding activity.

An EmaA homolog of the closely related species *Aggregatibacter aphrophilus* (formerly *Haemophilus aphrophilus*) is reported to be glycosylated (60). This homolog shares ~30-34% amino acid identity with *A. actinomycetemcomitans* EmaA (http://blast.ncbi.nlm.nih.gov/Blast.cgi) but shares many of the classic hallmarks of the adhesin: an amino-terminal signal sequence, a passenger domain consisting of the functional head domain and a long coiled-coil stalk, and a carboxyl-terminal translocation membrane anchor domain (61–64). The head domain retains multiple (>14) conserved SVAIG-like sequences characteristic of a YadA-like type V_c_ autotransporter (61, 65, 66) but has a low amino acid sequence similarity (~27%) when compared with *A. actinomycetemcomitans* EmaA. Rempe *et al*. (60) reported that *A. aphrophilus* EmaA was N-glycosylated by an orthologue of the HMW1C glycosyltransferase first identified in *Haemophilus influenzae*. Located in the cytoplasm, HMW1C transfers glucose and galactose to asparagine residues within the high molecular weight protein (HMW1) of *H. influenzae*. These adhesins belong to the family of type V_b_ secretory proteins that consist of partnered subunits: the functional, extracellularly exposed domain (A) and the β-barrel translocation/membrane anchor domain (B) (67–69). Similar to EmaA, glycosylation of HMW1 is required for protein function and stability (70–72). Although proteins similar to HMW1C are prevalent among a diverse set of Gram negative bacterial families, including both the Pastuerellaceae and Enterobacteriaceae (67), no HMW1C analog appears to be present in the *A. actinomycetemcomitans* genome (data not shown, (60)).

The result of this study lends additional support to the hypothesis that WaaL plays a dual function in *A. actinomycetemcomitans* physiology (6, 7). The enzyme acts as a ligase to covalently attach O-PS to the core oligosaccharide of the growing LPS structure, as well as a participating in the modification of the adhesin required for collagen binding. This dual function is in stark contrast with the glycosylation of other prokaryotic proteins that are typically mediated by a dedicated glycosyltransferase (73, 74), some of which utilize glycans from the LPS biosynthetic pathway (75–78). However, it has been demonstrated that the glycosyltransferase PglB from *Campylobacter jejuni* can mediate the transfer of multiple different polysaccharides from the Und-PP carrier to an acceptor protein (79), which lends credence to the role of WaaL in EmaA modification.

The multiple roles for WaaL ligase appear to be a novel paradigm adapted by this oral bacterium. Additionally, the ability of the *H. influenzae* WaaL to complement collagen binding in the *A. actinomycetemcomitans* mutant strain suggests that this multi-functional glycosyltransferase may have similar, as yet unidentified role(s) in other bacterial species. The combination of physiological functions of this enzyme may have some evolutionary advantage for the virulence of *A. actinomycetemcomitans* in oral and systemic diseases.

## Acknowledgements

We would like to acknowledge the National BioResource Project (NBRP) of the National Institute of Genetics, Japan, for the *E. coli* strains JW3597-KC and ME9062. We would like to acknowledge the late Dr. Edward Lally at the University of Pennsylvania School of Dental Medicine for the hemolytic *E. coli* strain 362-1430 used in this study. We would like to thank Dr. Joseph St. Geme at the Children’s Hospital of Philadelphia for the *H. influenzae* strain Rd used in this study. We would like to thank Dr. Marcin Filutowicz at the University of Wisconsin-Madison for providing the plasmid pJD104 used in this study. We would like to thank Ulrich Eckhard at the Institute of Molecular Biology of Barcelona (IBMB-CSIC) for providing the ColH plasmid used in this study. We would also like to thank Teresa Ruiz for editorial assistance. This research was supported by grant 5R01DE024554 from National Institutes of Health/National Institutes for Dental Craniofacial Research (NIH/NIDCR).

## Author Contributions

David R. Danforth: Conceptualization, Methodology, Investigation, Writing (Original Draft, Review, and Editing).

Marcella Melloni, Richard Thorpe, Avi Cohen, Richard Voogt, Jake Tristano: Investigation, Writing (Review and Editing).

Keith P. Mintz: Conceptualization, Methodology, Writing (Original Draft, Review, and Editing), Supervision, Project Administration.

---

**Supplementary Fig. 1**. Functional activity of chimeric WaaL expressed *in trans* in an *A. actinomycetemcomitans waaL* mutant strain. Representative collagen binding assay. Strain VT1169 (wild type), isogenic *waaL* mutant (mutant), mutant transformed with a plasmid expressing *E. coli waaL* (*E. c*.), mutant transformed with a plasmid expressing the *E. coli*/*A. actinomycetemcomitans* chimeric *waaL* (chimera), mutant transformed with a plasmid expressing *A. actinomycetemcomitans waaL* (*A. a*.). Performed in triplicate. The statistical significance is indicated with an asterisk (*p* < 0.05).

**Supplementary Fig. 2**. Comparison of relative EmaA expression levels expressed *in trans* in *E. coli* and *A. actinomycetemcomitans*. Immuno-dot blot analysis. Outer membrane fractions were quantified, and similar concentrations of protein were loaded into the wells before being probed using a monoclonal antibody specific for EmaA. *E. coli* strain MG1655 (*E. coli*), MG1655 transformed with a plasmid expressing *emaA* (*E. coli: emaA^+^*); *A. actinomycetemcomitans emaA* isogenic mutant strain KM73 (*A. a*.), KM73 transformed with a plasmid expressing *emaA* (*A. a.: emaA^+^*).

**Supplementary Fig. 3**. Collagen binding activity of *emaA* modified with a collagenase cut site in an *emaA* mutant *A. actinomycetemcomitans* strain. Strain KM73 (mutant), KM73 transformed with a plasmid expressing *emaA* (*emaA^+^*), KM73 transformed with a plasmid expressing collagenase cut site modified *emaA* (modified *emaA^+^*). Representative assay, performed in triplicate. The statistical significance is indicated with an asterisk (*p* < 0.05).

## References

1. Silhavy TJ, Kahne D, Walker S. 2010. The bacterial cell envelope. Cold Spring Harb Perspect Biol 2:a000414.

2. Beveridge TJ. 1999. Structures of gram-negative cell walls and their derived membrane vesicles. J Bacteriol 181:4725–33.

3. Bertani B, Ruiz N. 2018. Function and Biogenesis of Lipopolysaccharides. EcoSal Plus 8.

4. Raetz CR, Whitfield C. 2002. Lipopolysaccharide endotoxins. Annu Rev Biochem 71:635–700.

5. Whitfield C, Amor P, Köplin R. 1997. Modulation of the surface architecture of gram-negative bacteria by the action of surface polymer:lipid A-core ligase and by determinants of polymer chain length. Mol Microbiol 23:629–38.

6. Tang G, Mintz KP. 2010. Glycosylation of the collagen adhesin EmaA of *Aggregatibacter actinomycetemcomitans* is dependent upon the lipopolysaccharide biosynthetic pathway. J Bacteriol 192:1395–404.

7. Tang G, Ruiz T, Mintz KP. 2012. O-polysaccharide glycosylation is required for stability and function of the collagen adhesin EmaA of *Aggregatibacter actinomycetemcomitans*. Infect Immun 80:2868–77.

8. Darveau RP. 2010. Periodontitis: a polymicrobial disruption of host homeostasis. Nat Rev Microbiol 8:481–90.

9. Zambon J, Slots J, Genco R. 1983. Serology of oral *Actinobacillus actinomycetemcomitans* and serotype distribution in human periodontal disease. Infect Immun 41:19–27.

10. Fine DH, Patil AG, Velusamy SK. 2019. *Aggregatibacter actinomycetemcomitans* (*Aa*) Under the Radar: Myths and Misunderstandings of Aa and Its Role in Aggressive Periodontitis. Front Immunol 10:728.

11. Fine DH, Markowitz K, Fairlie K, Tischio-Bereski D, Ferrendiz J, Furgang D, Paster BJ, Dewhirst FE. 2013. A consortium of *Aggregatibacter actinomycetemcomitans*, *Streptococcus parasanguinis*, and *Filifactor alocis* is present in sites prior to bone loss in a longitudinal study of localized aggressive periodontitis. J Clin Microbiol 51:2850–61.

12. Zambon JJ. 1985. *Actinobacillus actinomycetemcomitans* in human periodontal disease. J Clin Periodontol 12:1–20.

13. Slots J, Ting M. 2000. *Actinobacillus actinomycetemcomitans* and *Porphyromonas gingivalis* in human periodontal disease: occurrence and treatment. Periodontol 2000 20:82–121.

14. Haubek D, Ennibi O, Poulsen K, Vaeth M, Poulsen S, Kilian M. 2008. Risk of aggressive periodontitis in adolescent carriers of the JP2 clone of *Aggregatibacter (Actinobacillus) actinomycetemcomitans* in Morocco: a prospective longitudinal cohort study. Lancet 371:237–42.

15. Kaplan AH, Weber DJ, Oddone EZ, Perfect JR. 1989. Infection due to *Actinobacillus actinomycetemcomitans*: 15 cases and review. Rev Infect Dis 11:46–63.

16. van Winkelhoff AJ, Slots J. 1999. *Actinobacillus actinomycetemcomitans* and *Porphyromonas gingivalis* in nonoral infections. Periodontology 2000 20:122–135.

17. Montemurro N, Perrini P, Marani W, Chaurasia B, Corsalini M, Scarano A, Rapone B. 2021. Multiple Brain Abscesses of Odontogenic Origin. May Oral Microbiota Affect Their Development? A Review of the Current Literature. Applied Sciences 11:3316.

18. Berbari EF, Cockerill FR, 3rd, Steckelberg JM. 1997. Infective endocarditis due to unusual or fastidious microorganisms. Mayo Clin Proc 72:532–42.

19. Goldberg MH, Katz J. 2006. Infective endocarditis caused by fastidious oro-pharyngeal HACEK micro-organisms. J Oral Maxillofac Surg 64:969–71.

20. Das M, Badley AD, Cockerill FR, Steckelberg JM, Wilson WR. 1997. Infective endocarditis caused by HACEK microorganisms. Annu Rev Med 48:25–33.

21. Paturel L, Casalta JP, Habib G, Nezri M, Raoult D. 2004. *Actinobacillus actinomycetemcomitans* endocarditis. Clin Microbiol Infect 10:98–118.

22. Tang G, Kitten T, Munro CL, Wellman GC, Mintz KP. 2008. EmaA, a potential virulence determinant of *Aggregatibacter actinomycetemcomitans* in infective endocarditis. Infect Immun 76:2316–24.

23. Ruiz T, Lenox C, Radermacher M, Mintz KP. 2006. Novel surface structures are associated with the adhesion of *Actinobacillus actinomycetemcomitans* to collagen. Infect Immun 74:6163–70.

24. Yu C, Mintz KP, Ruiz T. 2009. Investigation of the three-dimensional architecture of the collagen adhesin EmaA of *Aggregatibacter actinomycetemcomitans* by electron tomography. J Bacteriol 191:6253–61.

25. Tang G, Ruiz T, Barrantes-Reynolds R, Mintz KP. 2007. Molecular heterogeneity of EmaA, an oligomeric autotransporter adhesin of *Aggregatibacter* (*Actinobacillus*) *actinomycetemcomitans*. Microbiology 153:2447–2457.

26. Danforth DR, Tang-Siegel G, Ruiz T, Mintz KP. 2019. A Nonfimbrial Adhesin of *Aggregatibacter actinomycetemcomitans* Mediates Biofilm Biogenesis. Infect Immun 87.

27. Tang-Siegel GG, Danforth DR, Tristano J, Ruiz T, Mintz KP. 2022. The serotype a-EmaA adhesin of Aggregatibacter actinomycetemcomitans does not require O-PS synthesis for collagen binding activity. Microbiology (Reading) 168.

28. Watson A, Naughton H, Radermacher M, Mintz KP, Ruiz T. 2015. Tomographic Analysis of EmaA Adhesin Glycosylation in *Aggregatibacter actinomycetemcomitans*. Microscopy and Microanalysis 21:899–900.

29. Watson A, Tang-Siegel G, Brooks CJ, Radermacher M, Mintz KP, Ruiz T. 2016. Structural Significance of EmaA Glycosylation in *A. actinomycetemcomitans*. Microscopy and Microanalysis 22:1132–1133.

30. Kelley LA, Mezulis S, Yates CM, Wass MN, Sternberg MJE. 2015. The Phyre2 web portal for protein modeling, prediction and analysis. Nature Protocols 10:845–858.

31. Yang J, Zhang Y. 2015. I-TASSER server: new development for protein structure and function predictions. Nucleic Acids Res 43:W174–81.

32. Xu J, McPartlon M, Li J. 2021. Improved protein structure prediction by deep learning irrespective of co-evolution information. Nat Mach Intell 3:601–609.

33. Wang S, Ma J, Peng J, Xu J. 2013. Protein structure alignment beyond spatial proximity. Scientific Reports 3:1448.

34. Wang S, Peng J, Xu J. 2011. Alignment of distantly related protein structures: algorithm, bound and implications to homology modeling. Bioinformatics 27:2537–2545.

35. Mintz KP. 2004. Identification of an extracellular matrix protein adhesin, EmaA, which mediates the adhesion of *Actinobacillus actinomycetemcomitans* to collagen. Microbiology 150:2677–2688.

36. Rose JE, Meyer DH, Fives-Taylor PM. 2003. Aae, an autotransporter involved in adhesion of *Actinobacillus actinomycetemcomitans* to epithelial cells. Infect Immun 71:2384–93.

37. Mintz KP, Fives-Taylor PM. 1994. Adhesion of *Actinobacillus actinomycetemcomitans* to a human oral cell line. Infect Immun 62:3672–8.

38. Merritt JH, Kadouri DE, O’Toole GA. 2005. Growing and analyzing static biofilms. Curr Protoc Microbiol Chapter 1:Unit 1B 1.

39. Smith KP, Fields JG, Voogt RD, Deng B, Lam YW, Mintz KP. 2015. Alteration in abundance of specific membrane proteins of *Aggregatibacter actinomycetemcomitans* is attributed to deletion of the inner membrane protein MorC. Proteomics 15:1859–67.

40. Ducka P, Eckhard U, Schonauer E, Kofler S, Gottschalk G, Brandstetter H, Nuss D. 2009. A universal strategy for high-yield production of soluble and functional clostridial collagenases in *E. coli*. Appl Microbiol Biotechnol 83:1055–65.

41. Davis MR, Jr., Goldberg JB. 2012. Purification and visualization of lipopolysaccharide from Gram-negative bacteria by hot aqueous-phenol extraction. J Vis Exp doi:10.3791/3916.

42. Pérez J, McGarry M, Marolda C, Valvano M. 2008. Functional analysis of the large periplasmic loop of the *Escherichia coli* K-12 WaaL O-antigen ligase. Mol Microbiol 70:1424–40.

43. Ruan X, Loyola DE, Marolda CL, Perez-Donoso JM, Valvano MA. 2012. The WaaL O-antigen lipopolysaccharide ligase has features in common with metal ion-independent inverting glycosyltransferases. Glycobiology 22:288–99.

44. Ruan X, Monjarás Feria J, Hamad M, Valvano MA. 2018. *Escherichia coli* and *Pseudomonas aeruginosa* lipopolysaccharide O-antigen ligases share similar membrane topology and biochemical properties. Molecular Microbiology 110:95–113.

45. Datsenko KA, Wanner BL. 2000. One-step inactivation of chromosomal genes in *Escherichia coli* K-12 using PCR products. Proc Natl Acad Sci U S A 97:6640–5.

46. Baba T, Ara T, Hasegawa M, Takai Y, Okumura Y, Baba M, Datsenko KA, Tomita M, Wanner BL, Mori H. 2006. Construction of *Escherichia coli* K-12 in-frame, single-gene knockout mutants: the Keio collection. Mol Syst Biol 2:2006.0008.

47. Azari F, Radermacher M, Mintz K, Ruiz T. 2012. Correlation of the amino-acid sequence and the 3D structure of the functional domain of EmaA from *Aggregatibacter actinomycetemcomitans*. J Struct Biol 177:439–46.

48. Danforth DR, Melloni M, Tristano J, Mintz KP. 2021. Contribution of adhesion proteins to *Aggregatibacter actinomycetemcomitans* biofilm formation. Mol Oral Microbiol 36:243–253.

49. Valvano MA. 2011. Common themes in glycoconjugate assembly using the biogenesis of O-antigen lipopolysaccharide as a model system. Biochemistry (Mosc) 76:729–35.

50. Abeyrathne P, Daniels C, Poon K, Matewish M, Lam J. 2005. Functional Characterization of WaaL, a Ligase Associated with Linking O-Antigen Polysaccharide to the Core of *Pseudomonas aeruginosa* Lipopolysaccharide. J Bacteriol 187:3002–3012.

51. Schild S, Lamprecht AK, Reidl J. 2005. Molecular and functional characterization of O antigen transfer in *Vibrio cholerae*. J Biol Chem 280:25936–47.

52. Abeyrathne P, Lam J. 2007. WaaL of *Pseudomonas aeruginosa* utilizes ATP in in vitro ligation of O antigen onto lipid A-core. Mol Microbiol 65:1345–59.

53. Heinrichs DE, Monteiro MA, Perry MB, Whitfield C. 1998. The assembly system for the lipopolysaccharide R2 core-type of *Escherichia coli* is a hybrid of those found in *Escherichia coli* K-12 and *Salmonella enterica*. Structure and function of the R2 WaaK and WaaL homologs. J Biol Chem 273:8849–59.

54. Heinrichs DE, Yethon JA, Whitfield C. 1998. Molecular basis for structural diversity in the core regions of the lipopolysaccharides of *Escherichia coli* and *Salmonella enterica*. Molecular Microbiology 30:221–232.

55. Han W, Wu B, Li L, Zhao G, Woodward R, Pettit N, Cai L, Thon V, Wang PG. 2012. Defining function of lipopolysaccharide O-antigen ligase WaaL using chemoenzymatically synthesized substrates. J Biol Chem 287:5357–65.

56. Stenutz R, Weintraub A, Widmalm G. 2006. The structures of *Escherichia coli* O-polysaccharide antigens. FEMS Microbiology Reviews 30:382–403.

57. Perry MB, MacLean LL, Gmur R, Wilson ME. 1996. Characterization of the O-polysaccharide structure of lipopolysaccharide from *Actinobacillus actinomycetemcomitans* serotype b. Infect Immun 64:1215–9.

58. Liu D, Reeves PR. 1994. *Escherichia coli* K12 regains its O antigen. Microbiology 140:49–57.

59. Tang-Siegel GG, Radermacher M, Mintz KP, Ruiz T. 2022. Serotype specific sugars impact structure but not functions of the trimeric autotransporter adhesin EmaA of *Aggregatibacter actinomycetemcomitans*. J Bacteriol In press.

60. Rempe KA, Spruce LA, Porsch EA, Seeholzer SH, Nørskov-Lauritsen N, St Geme JW, 3rd. 2015. Unconventional N-Linked Glycosylation Promotes Trimeric Autotransporter Function in Kingella kingae and Aggregatibacter aphrophilus. mBio 6.

61. Yu C, Ruiz T, Lenox C, Mintz KP. 2008. Functional mapping of an oligomeric autotransporter adhesin of *Aggregatibacter actinomycetemcomitans*. J Bacteriol 190:3098–109.

62. Tamm A, Tarkkanen AM, Korhonen TK, Kuusela P, Toivanen P, Skurnik M. 1993. Hydrophobic domains affect the collagen-binding specificity and surface polymerization as well as the virulence potential of the YadA protein of *Yersinia enterocolitica*. Mol Microbiol 10:995–1011.

63. Mühlenkamp M, Oberhettinger P, Leo JC, Linke D, Schütz MS. 2015. *Yersinia* adhesin A (YadA)--beauty & beast. Int J Med Microbiol 305:252–8.

64. Roggenkamp A, Ackermann N, Jacobi CA, Truelzsch K, Hoffmann H, Heesemann J. 2003. Molecular analysis of transport and oligomerization of the *Yersinia enterocolitica* adhesin YadA. J Bacteriol 185:3735–44.

65. Nummelin H, Merckel MC, Leo JC, Lankinen H, Skurnik M, Goldman A. 2004. The Yersinia adhesin YadA collagen-binding domain structure is a novel left-handed parallel beta-roll. Embo J 23:701–11.

66. Tahir YE, Kuusela P, Skurnik M. 2000. Functional mapping of the *Yersinia enterocolitica* adhesin YadA. Identification Of eight NSVAIG - S motifs in the amino-terminal half of the protein involved in collagen binding. Mol Microbiol 37:192–206.

67. McCann JR, St Geme JW, 3rd. 2014. The HMW1C-like glycosyltransferases--an enzyme family with a sweet tooth for simple sugars. PLoS Pathog 10:e1003977.

68. Buscher AZ, Burmeister K, Barenkamp SJ, St Geme JW, 3rd. 2004. Evolutionary and functional relationships among the nontypeable Haemophilus influenzae HMW family of adhesins. J Bacteriol 186:4209–17.

69. Barenkamp SJ, St Geme JW, 3rd. 1994. Genes encoding high-molecular-weight adhesion proteins of nontypeable Haemophilus influenzae are part of gene clusters. Infect Immun 62:3320–8.

70. Grass S, Buscher AZ, Swords WE, Apicella MA, Barenkamp SJ, Ozchlewski N, St Geme JW, 3rd. 2003. The *Haemophilus influenzae* HMW1 adhesin is glycosylated in a process that requires HMW1C and phosphoglucomutase, an enzyme involved in lipooligosaccharide biosynthesis. Mol Microbiol 48:737–51.

71. Grass S, Lichti CF, Townsend RR, Gross J, St Geme JW, 3rd. 2010. The *Haemophilus influenzae* HMW1C protein is a glycosyltransferase that transfers hexose residues to asparagine sites in the HMW1 adhesin. PLoS Pathog 6:e1000919.

72. Gross J, Grass S, Davis A, Gilmore-Erdmann P, Townsend R, St Geme Jr. 2008. The Haemophilus influenzae HMW1 adhesin is a glycoprotein with an unusual N-linked carbohydrate modification. J Biol Chem 283:26010.

73. Brimer CD, Montie TC. 1998. Cloning and comparison of *fliC* genes and identification of glycosylation in the flagellin of *Pseudomonas aeruginosa* a-type strains. J Bacteriol 180:3209–17.

74. Szymanski CM, Yao R, Ewing CP, Trust TJ, Guerry P. 1999. Evidence for a system of general protein glycosylation in *Campylobacter jejuni*. Mol Microbiol 32:1022–30.

75. Lu Q, Yao Q, Xu Y, Li L, Li S, Liu Y, Gao W, Niu M, Sharon M, Ben-Nissan G, Zamyatina A, Liu X, Chen S, Shao F. 2014. An Iron-Containing Dodecameric Heptosyltransferase Family Modifies Bacterial Autotransporters in Pathogenesis. Cell Host & Microbe 16:351–363.

76. Moormann C, Benz I, Schmidt MA. 2002. Functional substitution of the TibC protein of enterotoxigenic *Escherichia coli* strains for the autotransporter adhesin heptosyltransferase of the AIDA system. Infect Immun 70:2264–70.

77. Yao Q, Lu Q, Wan X, Song F, Xu Y, Hu M, Zamyatina A, Liu X, Huang N, Zhu P, Shao F. 2014. A structural mechanism for bacterial autotransporter glycosylation by a dodecameric heptosyltransferase family. eLife 3:e03714.

78. Benz I, Schmidt MA. 2001. Glycosylation with heptose residues mediated by the *aah* gene product is essential for adherence of the AIDA-I adhesin. Mol Microbiol 40:1403–13.

79. Feldman MF, Wacker M, Hernandez M, Hitchen PG, Marolda CL, Kowarik M, Morris HR, Dell A, Valvano MA, Aebi M. 2005. Engineering N-linked protein glycosylation with diverse O antigen lipopolysaccharide structures in *Escherichia coli*. Proceedings of the National Academy of Sciences of the United States of America 102:3016–3021.

80. Mintz KP, Brissette C, Fives-Taylor PM. 2002. A recombinase A-deficient strain of *Actinobacillus actinomycetemcomitans* constructed by insertional mutagenesis using a mobilizable plasmid. Fems Microbiology Letters 206:87–92.

81. Sreenivasan PK, Fives-Taylor P. 1994. Isolation and characterization of deletion derivatives of pDL282, an *Actinobacillus actinomycetemcomitans/Escherichia coli* shuttle plasmid. Plasmid 31:207–14.

82. Eckhard U, Schonauer E, Brandstetter H. 2013. Structural basis for activity regulation and substrate preference of clostridial collagenases G, H, and T. J Biol Chem 288:20184–94.

