## Supplementary Fig for "Dual function of the O-antigen WaaL ligase of *Aggregatibacter actinomycetemcomitans*"

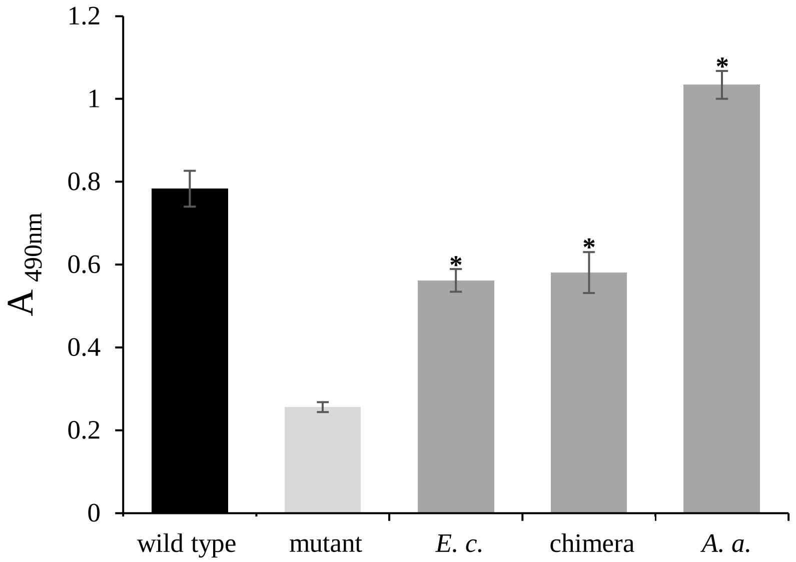


**Supplementary Fig. 1.** Functional activity of chimeric WaaL expressed *in trans* in an *A. actinomycetemcomitans waaL* mutant strain. Representative collagen binding assay. Strain VT1169 (wild type), isogenic *waaL* mutant (mutant), mutant transformed with a plasmid expressing *E. coli waaL* (*E. c.*), mutant transformed with a plasmid expressing the *E. coli*/*A. actinomycetemcomitans* chimeric *waaL* (chimera), mutant transformed with a plasmid expressing *A. actinomycetemcomitans waaL* (*A. a.*). Performed in triplicate. The statistical significance is indicated with an asterisk ($p < 0.05$).

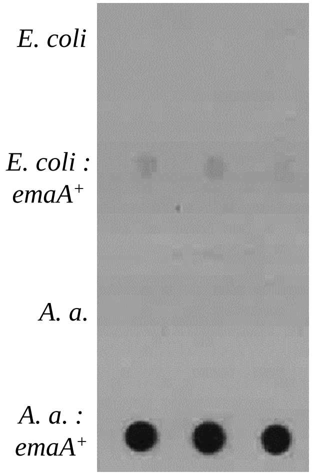


**Supplementary Fig. 2.** Comparison of relative EmaA expression levels expressed *in trans* in *E. coli* and *A. actinomycetemcomitans*. Immuno-dot blot analysis. Outer membrane fractions were quantified, and similar concentrations of protein were loaded into the wells before being probed using a monoclonal antibody specific for EmaA. *E. coli* strain MG1655 (*E. coli*), MG1655 transformed with a plasmid expressing *emaA* (*E. coli : emaA*$^{+}$); *A. actinomycetemcomitans emaA* isogenic mutant strain KM73 (*A. a.*), KM73 transformed with a plasmid expressing *emaA* (*A. a. : emaA*$^{+}$).

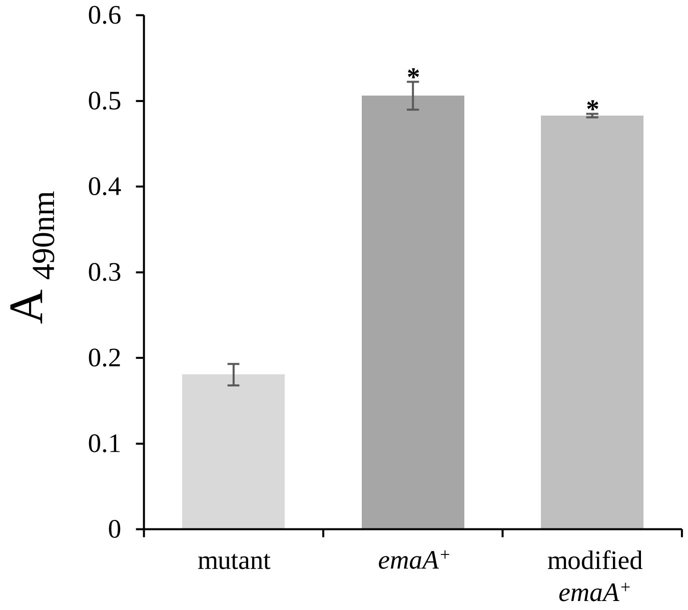


**Supplementary Fig. 3.** Collagen binding activity of *emaA* modified with a collagenase cut site in an *emaA* mutant *A. actinomycetemcomitans* strain. Strain KM73 (mutant), KM73 transformed with a plasmid expressing *emaA* (*emaA*$^+$), KM73 transformed with a plasmid expressing collagenase cut site modified *emaA* (modified *emaA*$^+$). Representative assay, performed in triplicate. The statistical significance is indicated with an asterisk ($p < 0.05$).
